# Evidence of backcross inviability and mitochondrial DNA paternal leakage in sea turtle hybrids

**DOI:** 10.1101/2022.10.10.511144

**Authors:** Sibelle T. Vilaça, Francesco Maroso, Paulo Lara, Benoit de Thoisy, Damien Chevallier, Larissa Souza Arantes, Fabricio R. Santos, Giorgio Bertorelle, Camila J. Mazzoni

## Abstract

Hybridization is known to be part of many species’ evolutionary history. Sea turtles have a fascinating hybridization system in which species separated by as much as 43 million years are still capable of hybridizing. Indeed, the largest nesting populations in Brazil of loggerheads (*Caretta caretta*) and hawksbills (*Eretmochelys imbricata*) have a high incidence of hybrids between these two species. A third species, olive ridleys (*Lepidochelys olivacea*), is also known to hybridize although at a smaller scale. Here we used restriction site-associated DNA sequencing (RAD-Seq) markers, mitogenomes, and satellite-telemetry to investigate the patterns of hybridization and introgression in the Brazilian sea turtle population and their relationship with the migratory behaviors between feeding and nesting aggregations. We also explicitly test if the mixing of two divergent genomes in sea turtle hybrids causes mitochondrial paternal leakage. We developed a new species-specific PCR-assay capable of detecting mitochondrial DNA (mtDNA) inheritance from both parental species and performed ultra-deep sequencing to estimate the abundance of each mtDNA type. Our results show that all adult hybrids are first generation (F1) and most display a loggerhead migratory behavior. We detected paternal leakage in F1 hybrids and different proportions of mitochondria from maternal and paternal species. Although previous studies showed no significant fitness decrease in hatchlings, our results support genetically-related hybrid breakdown possibly caused by cytonuclear incompatibility. Further research on hybrids from other populations besides Brazil and between different species will show if backcross inviability and mitochondrial paternal leakage is observed across sea turtle species.

## 1. INTRODUCTION

Interspecific hybridization is an important and fascinating evolutionary process in a species’ history (Noor & Bennett, 2009). The degree of hybridization and introgression, or repeated backcrossing of hybrids, to one or both parental species in natural systems is limited by reproductive isolating barriers. Increasing evidence supports these as permeable filters to gene flow (Twyford & Ennos, 2012). Therefore, extensive gene flow may occur between species where their distributions overlap. It has been estimated that between 1 to 10% of animals hybridize with at least one species (Mallet, 2005), but low hybrid viability may limit introgression depending on the degree of genomic divergence between parental species (Edmands, 2002). Generally, recently diverged species will produce fertile hybrids more frequently than species separated by longer times (Edmands, 2002). However, exceptions exist especially in reptiles where species diverged tens of million years can still interbreed and produce fertile offspring (Karl, Bowen, & Avise, 1995; Prager & Wilson, 1975).

Sea turtles have been swimming in earth’s oceans since the Cretaceous period (∼100 million years ago). Two families are currently recognized: the Dermochelyidae, represented by one species, the leatherback *Dermochelys coriacea*, and the Cheloniidae, which includes the other six extant species. Cheloniidae species are known to frequently hybridize, despite their ancient initial divergence ∼43 million years ago (Karl et al., 1995; Lara-Ruiz, Lopez, Santos, & Soares, 2006; Vilaça et al., 2021). In particular, four Cheloniidae species (loggerhead *Caretta caretta*, hawksbill *Eretmochelys imbricata*, olive ridley *Lepidochelys olivacea*, and green *Chelonia mydas* sea turtles) are known to hybridize and generate viable offspring in Brazil (Karl et al., 1995; Lara-Ruiz et al., 2006; Soares et al., 2018, 2021, 2017; Vilaça et al., 2012). This process is fairly common when compared to other worldwide nesting areas, especially in a small coastal region that comprises the states of Bahia and Sergipe where 32-42% of nesting hawksbills have been identified as hybrids between hawksbills and loggerheads (Lara-Ruiz et al., 2006; Soares et al., 2018, 2017; Vilaça et al., 2012). Other less common hybrids in Bahia include those between loggerheads x olive ridleys, hawksbills x olive ridleys (Soares et al., 2018, 2021, 2017; Vilaça et al., 2012), and greens x loggerheads (Karl et al., 1995). Interestingly, most molecular studies involving morphologically-identified hawksbills found hybrids with loggerhead mitochondrial DNA (mtDNA), and very few hybrid individuals displayed hawksbill mtDNA (Bass et al., 1996; Brito et al., 2020; Lara-Ruiz et al., 2006; Proietti et al., 2014; Soares et al., 2018, 2017).

Despite the high frequency of hybrids in this Brazilian region, it is still not clear why this is a unique phenomenon there and why those hybrids are almost always the offspring of a female loggerhead and a male hawksbill. Vilaça et al., (2021) raised the possibility that this is a direct cause of population decline, combined with overlapping reproductive peaks. Hawksbills are rarer than loggerheads in the Brazilian coast; hawksbills are classified as Critically Endangered by the International Union for Conservation of Nature (IUCN) (Mortimer & Donnelly, 2008), while loggerheads are classified as Least Concern in the southwest Atlantic Ocean and globally as Vulnerable (Casale & Tucker, 2017). In the Praia do Forte region (Bahia state) where hybrids have been found in large frequency, the nesting seasons of hawksbills and loggerheads slightly overlap (Marcovaldi & Dei Marcovaldi, 1999). The overlap of reproductive seasons, the temporal encounter between loggerhead females and hawksbill males, and the higher abundance of loggerheads in the Brazilian coastline, are all factors that may contribute to the observed gender-biased hybridization.

Although hybridization between divergent species is generally thought of as maladaptive (Delph & Demuth, 2016), reports of adaptive introgression with exchange of genes between species leading to a significant fitness benefit are rapidly increasing (Edelman & Mallet, 2021). In sea turtles, hybrids do not seem to be at great disadvantage compared to parental species. Analysis of fitness and reproductive output from Brazilian populations with largest proportions of hawksbill x loggerhead and olive ridley x loggerhead hybrids showed that fitness of hybrid hatchlings and reproductive output of hybrids (integrated over 7-years) is not different from both parental species (Soares et al., 2018, 2021, 2017). Furthermore, fitness reduction in the generations after F1, the so-called hybrid breakdown effect, was not observed among hatchlings in the Praia do Forte population (Soares et al., 2018). The lack of evident hybrid breakdown or long-term fitness decrease in loggerhead x hawksbill hatchlings poses an interesting evolutionary question, as species that diverged millions of years ago (Vilaça et al., 2021) apparently have compatible genomes and hybrids with well adapted phenotypes. The combination of two divergent genomes in hybrids can favour the colonization of unique niches (Taylor & Larson, 2019), even though hybrids can also assume intermediate niches or behavioral traits of the two parental species (Vilaça, Lima, Mazzoni, Santos, & de Thoisy, 2019). An important characteristic of sea turtles is the migratory pattern between nesting and foraging sites. Both hawksbills and loggerheads travel long distances (e.g., 1400 to 2400 km within the Brazilian continental shelf (Marcovaldi, Lopez, Soares, & López-Mendilaharsu, 2012), but the feeding areas of each species along the Brazilian coast are distinct. Hybrids were found to migrate to foraging areas typical of both parental species (Marcovaldi et al., 2010, 2012). However, the relationship between the hybrids’ genetic composition and their migratory pattern is not clear since a combination of mtDNA sequences (Lara-Ruiz et al., 2006) and few nuclear loci identified them as first generation (F1) (Arantes, Vilaça, Mazzoni, & Santos, 2020; Vilaça et al., 2012). The limits of these studies concern both the use of mtDNA and the number of nuclear loci.

Mitochondrial loci are by far the most common molecular markers used in sea turtle studies (Bowen & Karl, 2007). The mtDNA is a double-stranded circular genome averaging about 16,000 bp and is predominantly inherited as a single locus though the maternal lineage (Ballard & Whitlock, 2004). Previous research showed that mitochondrial heteroplasmy, or the presence of different mtDNA copies within one cell or individual, is common in green sea turtles and plays an important role in their ability to maintain diversity during population bottlenecks and founder effects (Tikochinski, Carreras, Tikochinski, & Vilaça, 2020). Although the mtDNA is regarded as having strict maternal inheritance in animals (Ballard & Whitlock, 2004), paternal leakage, the transmission of the paternal mitochondria to offspring, has been shown to occur in various species and is often associated with inter-specific hybridization (Fontaine, Cooley, & Simon, 2007; Kmiec, Woloszynska, & Janska, 2006) and also with intra-species crossings (Konrad et al., 2017; Kvist, Martens, Nazarenko, & Orell, 2003). Hybridization between divergent species was hypothesized to promote the breakdown of mechanisms that maintain strict maternal inheritance, like cytonuclear controls that limit the persistence of paternally-derived mitochondria (Breton & Stewart, 2015). Although the precise cellular mechanism responsible for enforcing mitochondrial maternal inheritance is unknown, hybridization between extremely divergent species, like sea turtles, poses an interesting model to test if paternal leakage can indeed be observed as a result of mixing two divergent genomes.

Although thousands of sea turtle individuals have been sequenced so far in different studies, the low number of molecular markers did not provide enough resolution to distinguish hybrid generations past F1 in the biggest hybrid population known so far (Soares et al., 2018; Vilaça et al., 2012). Therefore, no studies have provided enough resolution to investigate if the largest Brazilian nesting population is a hybrid swarm (when multiple hybrid generations due to a high proportion of backcross with one parental species or another hybrid), or if it is mostly formed by pure individuals and F1 hybrids. Furthermore, a better characterization of this population may enlighten behavioral patterns such as migration, and possibly improve conservation measures. Genome-wide markers, such as loci from restriction sites with associated DNA sequencing (RAD-seq, Davey et al., 2011), have been widely used in non-model species and could provide important genome insights by providing higher resolution in identifying later hybrid generations.

In this study, we investigated the hybridization process between hawksbill and loggerhead sea turtles from different nesting and feeding populations in the Brazilian coast using genomic markers. We were specifically interested in the introgression pattern in the Praia do Forte rookery, Bahia, the relationship between hybridization and migratory behavior, and mtDNA paternal leakage in sea turtle hybrids. We used the double digest RADseq (ddRAD) technique to genotype individuals with a large panel of Single Nucleotide Polymorphisms (SNPs), and used these genotypes to identify the genomic composition of hybrids and putative pure individuals. We also used telemetry data combined with hybrid generation assignment to infer if migratory patterns can be explained by introgression. To infer paternal leakage, we first sequenced mitogenomes for multiple individuals and then developed a targeted species-specific PCR test capable of detecting mtDNA inheritance from both species, combined with mtDNA ultra-deep sequencing to infer abundance of each mtDNA type.

## 2 METHODS

### 2.1. Sampling

We obtained a total of 129 tissue samples from Brazilian rookery and feeding populations (Figure 1). Samples of 41 individuals (loggerhead = 24, hawksbill = 17) from three Brazilian nesting sites (states of Bahia, Espírito do Santo, and Rio Grande do Norte, Supplementary Material Table S1) were used for this study. Samples were first sequenced for their D-loop region of the mitochondrial DNA following Vilaça, Lara-Ruiz, Marcovaldi, Soares, & Santos, (2013), which is usually used as a first identification of hybrids. In addition, we analysed 46 additional samples (loggerheads = 14, hawksbills = 20, hybrid = 12) that were used in a previous study based on 5 nuclear markers and with a previously known hybrid/pure status (Vilaça et al., 2012). All samples were collected by trained members of Fundação Projeto TAMAR. Samples were always identified as a parental species, never as hybrids. To identify potential hybrids with other sea turtle species we also included 8 green sea turtles from Brazil, French Guiana, Guadeloupe, and Martinique; and 7 olive ridleys from Brazil and French Guiana.

**Figure 1.**
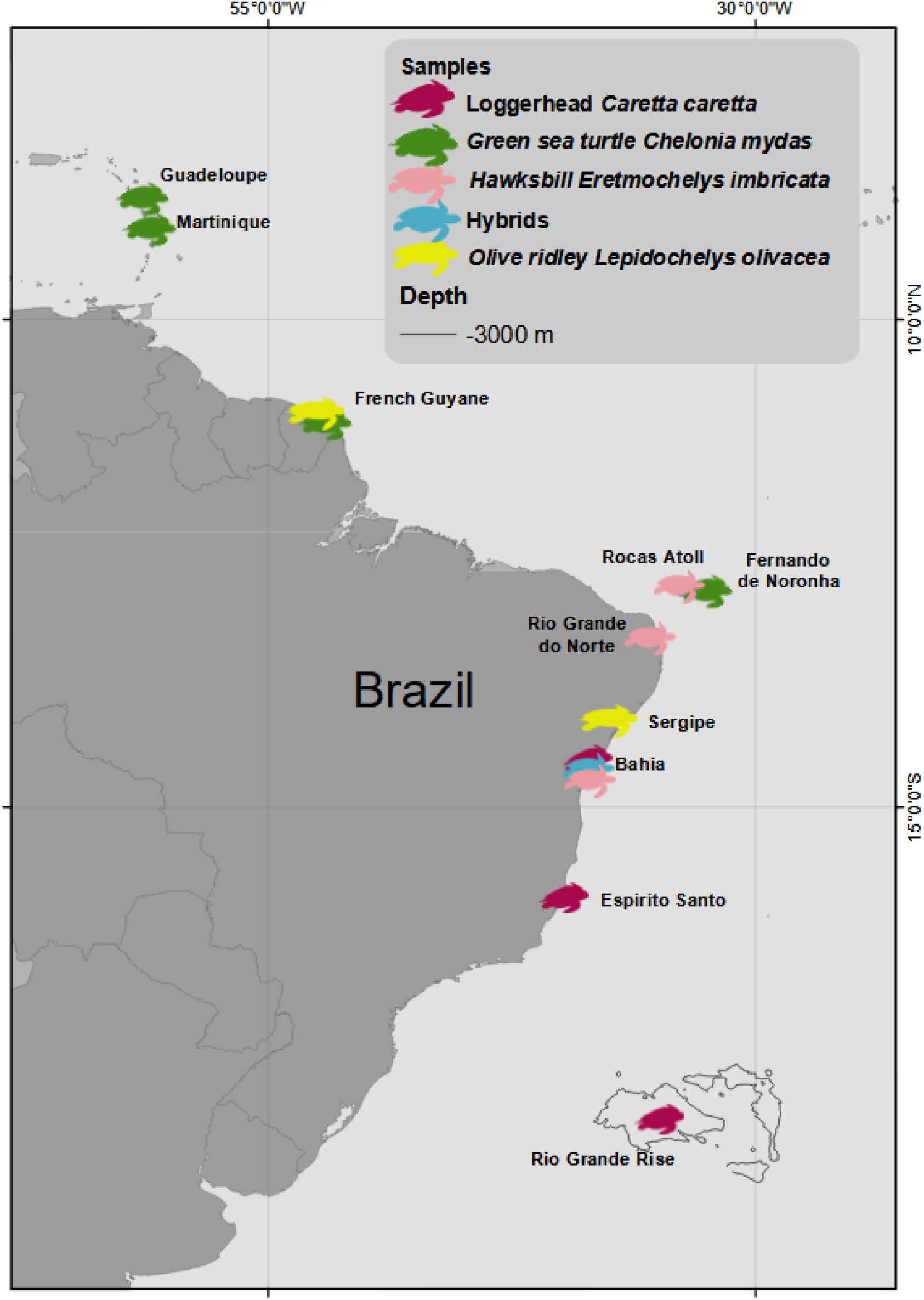
Map displaying sampling sites of samples used in this study.

### 2.2. ddRAD laboratory

DNA extractions were performed using the DNeasy Blood and Tissue kit (Qiagen). Double digest RAD (ddRAD) libraries were produced using the protocol of Peterson, Weber, Kay, Fisher, & Hoekstra, (2012) with modifications of adapter sequences according to Meyer & Kircher, (2010). In brief, 1 μg of genomic DNA was digested with two enzymes (EcoRI-HF and MseI) at 37 °C for at least two hours in a reaction volume dependent of the DNA concentration per sample. The adapter ligation step was performed immediately after the digestion. The P5 adapter had inline barcodes of varying sizes (5 to 9 bp) to avoid low diversity problems at the restriction site positions, and P7 adapter was designed with one strand lacking the indexing primer complementary region. Ligated samples were cleaned with 1.8X CleanPCR magnetic beads (CleanNA). A 10-cycle indexing PCR was performed independently for each individual, using one of the 50 indexes in the P7 adapter (Meyer & Kircher, 2010). The PCRs were cleaned with 0.8X CleanPCR magnetic beads, and concentration was measured with Qubit 2.0 using the dsDNA HS assay (Life Technologies). The indexed libraries were equimolarly pooled before the size selection step. The size selection step was performed with the Blue Pippin (Sage Science) using a 1.5% cassette and the R2 marker, targeting fragments between 490 and 610 bp. The final libraries were characterized with a qPCR using the KAPA SYBR FAST (Kapa Biosystems) kit and also checked in the Agilent 2100 Bioanalyzer High Sensitivity DNA (Agilent Technologies). The libraries were run in the in-house Illumina NextSeq 500 using the 300-cycle mid-or high-output kit.

### 2.3. ddRAD analysis

Raw reads quality was checked with FastQC (Andrews, 2010) and pre-processed for quality as described in Driller et al., (2020) with modifications to accommodate the lack of overlapping reads. Adapter sequences were trimmed using Cutadapt (Martin, 2011). P5 inline barcodes were demultiplexed using FLEXBAR v.3.0.3 (Roehr, Dieterich, & Reinert, 2017). Paired-end reads were merged using PEAR v.0.9.11 (Zhang, Kobert, Flouri, & Stamatakis, 2014) setting the maximum length of the assembled sequences to 249 bp to exclude short fragments. Unassembled R1 and R2 reads were trimmed to the maximum length of 140 and 135 bp, respectively, using Trimmomatic v.0.3.6 (Bolger, Lohse, & Usadel, 2014). The first five bases of R2 reads were also trimmed, as they had a below average Phred quality score (Q). Only sequences with overall Q>25 were analyzed. We used the checkRestrictionSites.py custom script to check the presence of the MseI enzyme at the beginning of reads R1 and discarded all the paired reads with incorrect restriction sites. We also checked for undigested and chimeric sequences and removed all reads containing intact MseI and EcoRI restriction sites.

After quality check, the processed reads were mapped to a set of ddRAD single copy orthologue loci (R2SCO-MseI-EcoRI-384-448) for hawksbills and loggerheads from Driller et al., (2020). Reads were mapped using bowtie2 v.2.3.4.1 (Langmead & Salzberg, 2012) with default parameters. The mapped reads were then processed using the reference-based pipeline in Stacks 2.2 (Catchen, Hohenlohe, Bassham, Amores, & Cresko, 2013). We used Stacks’ *populations* module to obtain loci present in at least one population and in a minimum of 70% of individuals. We also compared if SNPs and microhaplotypes had a similar resolution in defining clusters, since microhaplotypes from RAD loci might have higher power in population assignments (Baetscher, Clemento, Ng, Anderson, & Garza, 2018). The called SNPs were used to reconstruct haplotypes using the script Create_Haplotype_Structure.py, which was also used to filter loci based on their minimum coverage of 15 reads and the minimum number of populations covering a locus (popCOV = 4). To maximize the number of RAD loci and variants used to describe hybrids, we chose to analyse two datasets. The first, with more SNPs, had only hawksbills, loggerheads and their hybrids. In the second dataset, samples of olive ridleys, olive ridley x loggerhead hybrids, and green turtles were also included. Both datasets were filtered for SNPs with a minimum depth of 15X and a minimum allele frequency (MAF) of 0.05.

To identify introgression in individuals and putative population structure, we ran the model-based Structure v2.3.4 (Pritchard, Stephens, & Donnelly, 2000) without population/species *a priori* information, with admixed model and unlinked sites. The number of simulated clusters (K) ranged from 1 to 10, each repeated 10 times; 50,000 iterations as burn-in followed by 200,000 iterations. Results were parsed with Evanno’s method using the online tool Structure Harvester (Earl & vonHoldt, 2012) to determine the most likely K. To evaluate the consistency of solutions to clustering (major and minor clusters) and summary results for each K across replicates, we used the online tool CLUMPAK (Kopelman, Mayzel, Jakobsson, Rosenberg, & Mayrose, 2015). We also ran a non-model-based clustering method to compare assignments with Structure results. Discriminant Analysis of Principal Components (Jombart, Devillard, & Balloux, 2010) was performed in the R (R Core Team, 2017) package *adegenet* (Jombart & Ahmed, 2011). The number of retained principal components (PC) was optimized with *xValDapc* method. In all cases a low number of PC (5-10) allowed for very good sample assignment and no overfitting. Two or three Discriminant Axis (DA) were retained depending on their eigenvalues. To estimate pairwise relatedness estimates among all individuals, we used vcftools v1.9 (Danecek et al., 2011) with the command –relatedness (Yang et al., 2010).

We used the NewHybrids (Anderson & Thompson, 2002) function *gi*.*nhybrids()* implemented in the R package dartR (Gruber, Unmack, Berry, & Georges, 2018) to assign each individual to a “hybrid category” (pure species, F1, F2 or backcross). We extracted SNPs that were fixed in loggerheads and hawksbills (Fst = 1) and had a minimum coverage of 15X. A dataset with loggerheads, hawksbills, and their hybrids was used to calculate hybrid category probability. Two hundred loci were selected by the program, either randomly or using average polymorphism information content (PIC) values based on the 200 most informative loci. To verify potential influences of setting prior populations, we ran NewHybrids without assigning pure individuals as priors, and also using pure individuals previously identified as non-hybrids by Structure and DAPC.

We used the D-statistics to calculate conflicting patterns between ancestral (“A” alleles) and derived (“B” alleles) to distinguish introgression from incomplete lineage sorting (ILS). The D statistic is based on a tree model of four populations, defined as (((P1,P2),P3),O), where P1, P2, and P3 are ingroups and O is the outgroup. Excesses in the “ABBA” or “BABA” patterns produce deviation from D = 0, supporting introgression between P2 and P3 over the null hypothesis of ILS. We used the package D-suite (Malinsky, Matschiner, & Svardal, 2020) and the command Dtrios. To identify potential origin of hybrids and admixture between sites, samples were divided as population of origin (parental species) based on genetic structure found in previous studies (Shamblin et al., 2014; Vilaça et al., 2013), or as hybrids. Loggerheads were divided in two nesting areas (Bahia and Espírito Santo) and one feeding (Rio Grande Elevation). Hawksbills were divided in two nesting areas (Bahia and Rio Grande do Norte) and one feeding (Rocas Atoll). Olive ridleys were considered a single population (Vilaça, Hahn, Naro-Maciel, et al., 2022). Green turtles were set as outgroup. Significant gene-flow was considered if Z-score > 3 and assessed using jackknifing. We also estimated the genome fraction involved in the admixture (Malinsky et al., 2020) by computing the *f*4-ratio using the same software.

### 2.4. Migratory behavior of hybrids

As some of the individuals used were included in previous telemetry studies, we evaluated if their migratory patterns could be related to the fraction of the genome assigned to each parental species. For example, if a hybrid displays a migratory behavior similar to a pure hawksbill, this could be caused by introgression/backcrossing with a hawksbill (F1 x hawksbill), as a larger proportion of its genome will have come from hawksbills. The same rationale is valid for a trajectory pattern similar to pure loggerheads. In the case of F1 hybrids, since they have 50% of each parental genome, we hypothesize they could either have a behavior similar to parental species if they a have similar niches/migratory trajectories as at least one of the parental species; or dissimilar to both parentals if they have a diverse or intermediate migratory behavior. We gathered data from Marcovaldi et al., (2010, 2012) and matched hybrid generation assignment (F1, backcross) from our Structure and NewHybrids assignments or from previous studies (Soares et al., 2021; Vilaça et al., 2012). For all analysed samples, we had genetic information to confirm their hybrid/pure status, even though not all were sequenced with genome-wide markers. Therefore, we obtained ddRAD data for 3 hybrids and 1 hawksbill. For 4 samples (3 hawksbills, 1 hybrid) genetic information was retrieved from Vilaça et al., (2012) for multiple nuclear loci. All loggerheads (n = 7) were previously analysed (Soares et al., 2021) and data for one mitochondrial and one nuclear locus was available. All individuals had their transmitters deployed at Praia do Forte (Bahia). Maps were done in ArcGIS 10 (ESRI) and minimum convex polygons and core foraging areas followed the methods from Marcovaldi et al., (2010, 2012).

### 2.5. Mitogenomes

We selected 37 samples (loggerheads = 16, hawksbills = 11, hybrids = 10) from three Brazilian rookeries and feeding sites. The only previous study with population-level mitogenomes of sea turtles showed low genetic diversity outside the D-loop region in Caribbean green sea turtles (Shamblin et al., 2012). Given that both loggerheads and hawksbills show generally lower genetic diversity in the mtDNA compared to green sea turtles (Jensen et al., 2019; Shamblin et al., 2014; Vilaça et al., 2013), we expect that our sampling number will recover most of the polymorphisms present in the Brazilian population. Five new primer combinations were developed using two published primers (Abreu-Grobois et al., 2006; Duchene et al., 2012) and six newly developed ones (Table S2). New primers based on previously published mitogenomes (Duchene et al., 2012) were designed for overlapping fragments longer than 3000 bp using Primer3 (Untergasser et al., 2012) implemented in Geneious 8.1.3 (Kearse et al., 2012). The amplification of four fragments consisted of a 40 μL mix containing 100-200 ng of genomic DNA, 1X Herculase II reaction buffer, 0.4 μL of dNTPs (at 25mM each), 1 μL of each primer at 10mM, 1% BSA, 1% DMSO and 0.8 μL of Herculase II Fusion DNA polymerase. The amplification reaction consisted of 95 °C for 3 minutes, followed by 35 cycles of 94 °C for 30 seconds, 55/57 °C for 30 seconds, 70 °C for 5 minutes and a final extension of 70 °C for 10 minutes. The fragments were checked in 1% agarose gel stained with Sybr gold. If any sample presented nonspecific amplification, the fragment of expected size was excised from the agarose gel and purified with the GeneJet PCR Purification kit (Thermo Scientific). For each sample, the four fragments were purified with 1.8X CleanPCR magnetic beads, had their concentration measured in Qubit 2.0 instrument applying the dsDNA assay (Life Technologies) and were equimolarly pooled. Each pool was sheared using a Covaris S220 ultrasonicator (Covaris) to a mean of 400 bp following the manufacturer’s recommendations. For the library construction we followed the protocol from Meyer & Kircher, (2010). The library was first size-selected in BluePippin using the 1.5% cassette with R2 marker (Sage Science) to a fragment size range between 490 bp to 610 bp. The final library was characterized with a qPCR using the KAPA SYBR FAST (Kapa Biosystems) kit and also checked with the Agilent Bioanalyzer 2100 High Sensitivity DNA chip. The sequencing was performed in the in-house NextSeq 500 with 2x 150 bp reads.

To assemble the mitogenomes we used the software Geneious 8.1.3. All sequences were imported into the software and mapped to two references (a hawksbill and a loggerhead) with haplotypes typical of the Atlantic Ocean (GenBank accession numbers JX454986 and JX454984). The mapping was done using the Geneious mapper with Custom sensitivity (minimum overlap identity: 98%, maximum gap size: 50, index word length: 12, allow gaps of maximum: 15%, minimum mapping quality: 30, maximum ambiguity: 4). Annotations were transferred from the references to the newly assembled mitogenomes. The mitogenomes were aligned with MUSCLE (Edgar, 2004) and gaps and polymorphisms were manually checked.

### 2.6. Paternal leakage detection from PCR

We developed a species-specific PCR-based test to detect if paternal leakage is present in sea turtle hybrids. We amplified a total of 118 samples, including those sequenced for ddRAD and previously sequenced for mtDNA and nuclear markers (Vilaça et al., 2013, 2012). We tested two olive ridley and green sea turtle samples to verify possible cross-amplification. Hybrids between other species or individuals with haplotypes characteristic of other oceans (CC-A2, CC-A33, CC-A34) present in Rio Grande Elevation feeding area were not included.

The full mitochondrial DNA of Brazilian loggerheads and hawksbills was used to find regions with enough differences to distinguish both species, and where primers could be designed. We used the same software as for the mitogenomes and searched for species-specific primers. We selected one primer pair per species with enough differences that would be species-specific (i.e., amplify one species but not in the other). The primers designed comprise a 2420 bp region between the 16S rRNA and ND2 loci. Primer sequences can be found in Table S3. Even though a smaller region could have been used, we preferred the use a longer region to avoid amplifying nuclear copies of mitochondrial DNA (NUMTs) (Richly & Leister, 2004). Sequences from non-common Atlantic haplotypes (common haplotypes include, e.g., CC-A4 for loggerheads and Ei8 for hawksbills) or from the Indo-Pacific Ocean might have mutations in the annealing sites, and since we did not have population-level mitogenomes for those haplotypes, we could not confirm if our primers would remain species-specific in individuals with mitochondrial haplotypes besides the ones sequenced here. Furthermore, in our amplification and sequencing efforts for mitogenomes, we observed that the same fragments failed to amplify in individuals with other (low frequency or non-Atlantic) haplotypes, which did not allow us to confirm the specificity of our primers in relation to other haplotypes.

Our species-specific PCR cocktail consisted of a 40 μL mix containing at least 8 ng of genomic DNA, 1X Phusion reaction buffer, 1 μL of dNTPs (at 10mM each), 1 μL of each primer at 10mM, and 0.25 μL of Phusion High-Fidelity DNA Polymerase (New England Biolabs). The amplification reaction consisted of 95 °C for 3 minutes, followed by 35 cycles of 94 °C for 30 seconds, 68 °C for 30 seconds, 70 °C for 1 minute and a final extension of 70 °C for 5 minutes. Each PCR included a blank, and a control sample for each species (negative and positive controls). The fragments were checked in 1% agarose gel stained with Sybr gold.

To perform detection tests in our primer pairs, we quantified the DNA of samples using the Qubit 2.0 instrument applying the dsDNA assay (Life Technologies). We chose one loggerhead and one hawksbill sample with the same amount of DNA (22 ng/μL). We tested the detection ability of each primer pair by serial diluting these samples (no dilution, 1:10, 1:100, 1:1000, 1:10,000) and amplifying them with their species-specific primer. The capacity of distinguishing both mitochondria was tested by mixing samples of both species in different concentrations (1:1, 1:10, 1:50, 1:100, 10:1, 50:1, 100:1) and amplifying them with the two species-specific primer pairs.

Our analysis consisted in observing the banding pattern for each sample. An individual was considered as having mtDNA paternal leakage only if it showed two bands of correct sizes in amplifications for both species. If a light band was detected, the PCR was repeated to confirm the presence of the second mitochondria. If it failed to show the two bands in the second PCR, the sample was considered as homoplasmic.

### 2.8. Paternal leakage detection from ultra-deep sequencing

To further confirm the presence in the hybrids of two different mitochondrial sequences and estimate the abundance of each mitochondrial type, we sequenced 40 samples using the Illumina platform. Samples were chosen as representatives of pure and hybrid categories from Brazilian populations that had enough good quality DNA. We used the primer pair corresponding to positions 9,518 and 12,629 (genes NADH dehydrogenase subunit 3 to NADH dehydrogenase subunit 5) (Table S2). This fragment was successfully amplified in all samples and did not overlap with the PCR-assay, providing further evidence that the entire mtDNA was inherited in individuals with evidence of paternal leakage. The PCR and library preparation followed the same protocol as for the mitogenome. To confirm the “leakage” pattern, 27 samples were sequenced in two different runs from independent PCRs and libraries. We subsampled 30,000 reads (expected mean coverage = 1000X) and mapped them to mitogenomes of Atlantic hawksbills (GenBank accession number: JX454986) and loggerheads (JX454984) using stringent mapping parameters so only sequences of its respective species were mapped to each reference. The mapping was done using the Geneious mapper with Custom sensitivity (minimum overlap identity: 98%, maximum gap size: 50, index word length: 12, allow gaps of maximum: 15%, minimum mapping quality: 30, maximum ambiguity: 4). To be considered as an indication of paternal leakage, an individual needed to show sequences with a homogeneous coverage of the entire amplified fragment and a minimum average coverage of 3 for both species’ mitochondrial DNA. Each individual was manually checked for mapping correctness to the right species and mapping patterns.

To infer the abundance of each parental mitochondria, we considered the ratio of sequences mapped to each species-specific mitogenome as a proxy for abundance. We divided the total number of sequences mapped to the less abundant mitochondrial reference divided by the number of sequences mapped to the most abundant mitochondria.

To further confirm the inheritance of mtDNA from both parents, we investigated if the second mitochondria might be derived from a NUMT. In humans, a previous study showed that biparental mtDNA transmission are cryptic mega-NUMTs that resemble paternal leakage (Wei et al., 2020). Thus, we aligned the hawksbill mtDNA to a draft assembly of the loggerhead genome (unpublished, N50 = 20.7Mb). Alignments were done using minimap2 v2.17 (Li, 2018) with maximum 5% of sequence divergence (asm5). We also checked the haplotypes obtained for all samples, as NUMTs tend to evolve slower than the mitochondrial counterparts. This means that NUMT sequences tend to be similar to the ancestral mitochondrial sequences, have random polymorphisms with respect to codon positions, and have also stop codons that disrupt the reading frame of protein-coding genes (Baldo, de Queiroz, Hedin, Hayashi, & Gatesy, 2011).

## 3. RESULTS

### 3.1. ddRAD and hybrid identification

For samples with high quality DNA (n = 95), we obtained a total of 255,738,252 sequences with an average coverage of 38.5X (details can be found in Table S1 and S4). Our dataset for loggerheads, hawksbills and their hybrids (n = 79) had a total of 2,138 loci and 7,845 variants. When including olive ridleys and green sea turtles (n = 95), we obtained a total of 11,740 variants in 1,368 RAD loci. Details for each processing step following Driller et al., (2020)’s protocol can be found in Table S4.

The Structure results subdivided hawksbills and loggerheads when K was set to two (Figure 2). All hybrids had a 50% component of each species, indicating that they are all first generation (F1) hybrids. The only exception is a hatchling previously known to be a F1 x hawksbill backcross. At K = 3, loggerheads from feeding areas with mtDNA haplotypes from Indo-Pacific (CC-A2, CC-A33, CC-A34) formed a different cluster, a pattern also seen in our relatedness analysis (Figure 3). When considering the four species, at K = 4 they formed separate homogeneous clusters (Figure 2C and 2D). One individual morphologically identified as a loggerhead and with an olive ridley mtDNA (haplotype Lo67, as defined in Vilaça et al., 2022) was shown to be an olive ridley x loggerhead F1 hybrid (OL). We did not observe any differences in cluster definition between SNPs and microhaplotypes (Figure 2).

**Figure 2.**
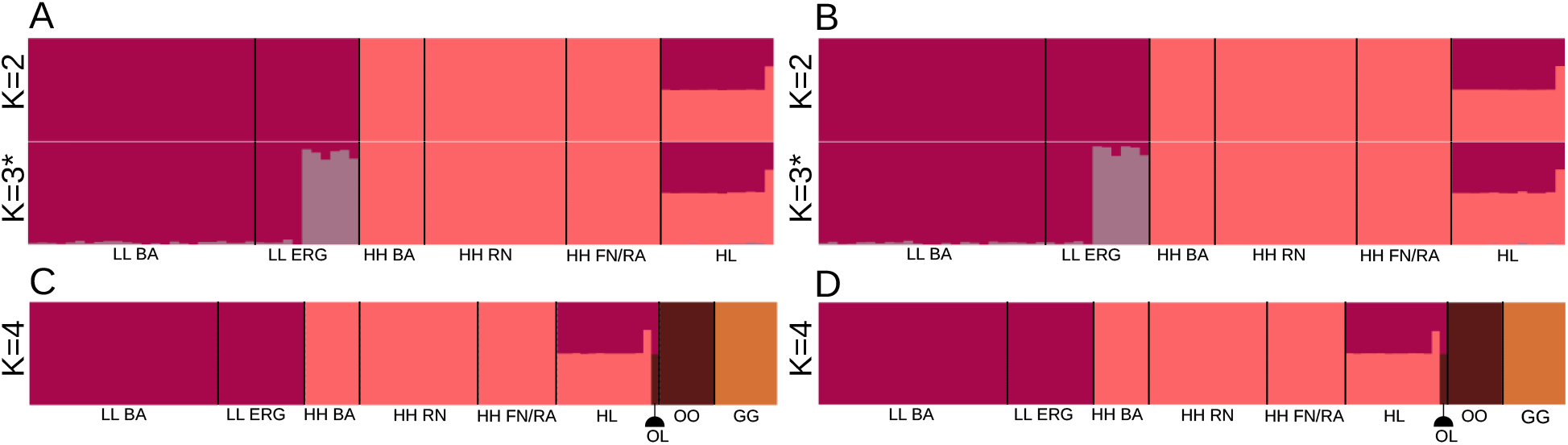
Proportional membership for hawksbills (HH), loggerheads (LL) and their hybrids (HL) using SNP (a) and haplotype (b) data; and including olive ridleys (OO), green sea turtles (GG) and their hybrids (OL), using SNP (c) and haplotype (d) information. Each individual is represented by a vertical bar, and the length of each bar indicates the probability of membership in each cluster.The asterisk denotes a minor cluster. Sampling sites are abbreviated as: BA= Bahia, ERG = Rio Grande elevation, RN = Rio Grande do Norte, FN = Fernando de Noronha, RA = Rocas Atoll.

**Figure 3.**
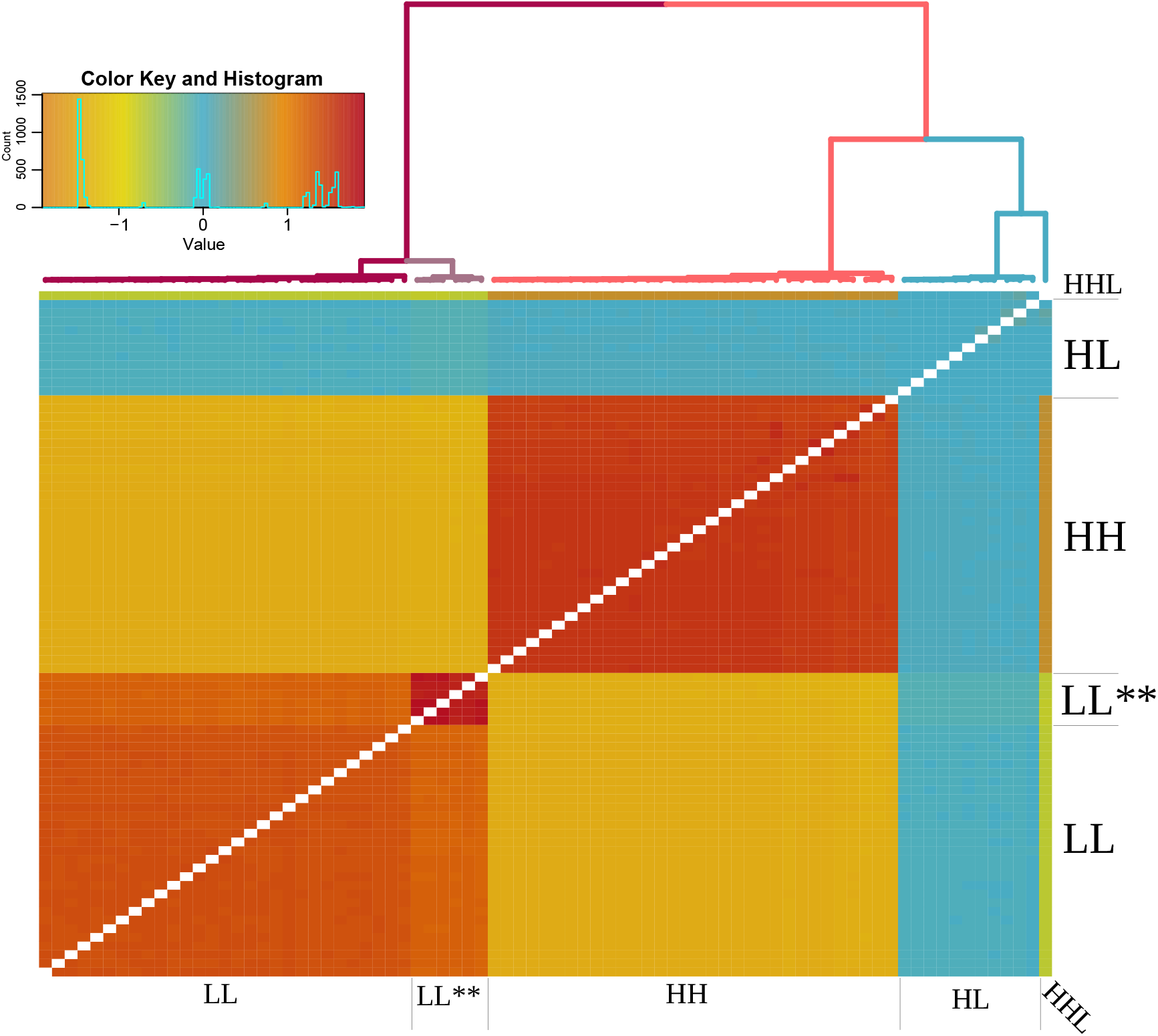
Pairwise relatedness analysis between hawksbills (HH), loggerheads (LL), F1 hybrids (HL) and backcrosses (HHL). LL** denotes the loggerhead individuals from the feeding population with Indo-Pacific and South African mtDNA haplotypes.

For the DAPC (Figures S1 and S2) and pairwise relatedness estimates (Figure 3), the results were similar to the Structure analysis, with hybrids clustering between loggerheads and hawksbills, with the exception of the backcross hatchling which is closer to hawksbills.

Individual assignments to a “hybrid category” with NewHybrids classified all hybrids as F1, with the exception of the backcrossed hatchling (Figure 4). This pattern was observed in all prior combinations.

**Figure 4.**
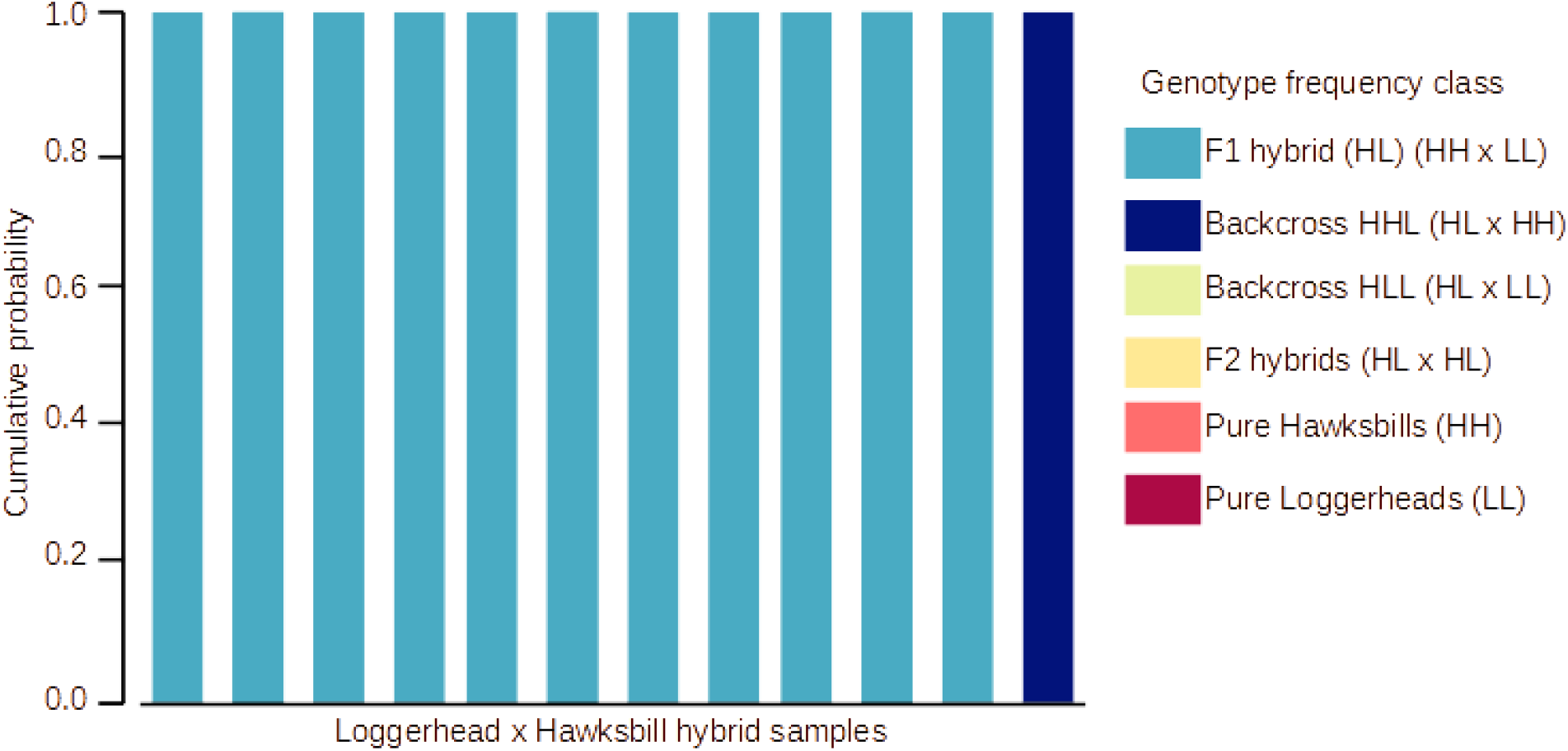
NewHybrids assignments. Only hybrid samples are shown, the remaining samples are all classified as pure hawksbills or loggerheads. The Y-axis is the cumulative posterior probability of each individual belonging to one of the six genotype frequency classes.

The role of incomplete lineage sorting in generating the observed hybridization patterns was excluded using D-statistics and *f*4-ratio (Table 1). First generation hawksbills x loggerheads hybrids and the single olive ridley x loggerhead hybrid had an estimate of admixture proportion (*f*4-ratio) expected for F1 hybrids (∼50%). The D-statistics, *f*4-ratio, and Z-scores were higher when considering trios when both P1 and P3 were individuals from Bahia (the same region as the hybrids), indicating that the hybridization is a local phenomenon of this population. For the backcrossed hatchling, the *f*4-ratio showed the expected combination of ∼25% of genome derived from loggerheads and 75% from hawksbills. No combinations between parental populations had a Z-score>3.

**Table 1.** D-statistics and *f*4-ratio results for hybrids and parental population combinations generated with D-suite. Since D-suite only calculates positive D values, only combinations which an excess of allele sharing between P2 and P3 populations are shown.

| P1 | P2 | P3 | D-stat | Z-score | p-<br>value | $f_4$-<br>ratio |
| --- | --- | --- | --- | --- | --- | --- |
| Hawksbill<br>Bahia | F1 hybrids | Loggerhead<br>Rookery | 0.980 | 347.9 | 0 | 0.501 |
| Hawksbill RN | F1 hybrids | Loggerhead<br>Rookery | 0.979 | 360.7 | 0 | 0.501 |
| Hawksbill<br>Feeding | F1 hybrids | Loggerhead<br>Rookery | 0.977 | 309.3 | 0 | 0.500 |
| Hawksbill<br>Bahia | F1 hybrids | Loggerhead<br>Feeding | 0.976 | 279.8 | 0 | 0.497 |
| Hawksbill RN | F1 hybrids | Loggerhead<br>Feeding | 0.976 | 286.8 | 0 | 0.497 |
| Hawksbill<br>Feeding | F1 hybrids | Loggerhead<br>Feeding | 0.973 | 259.7 | 0 | 0.497 |
| Hawksbill<br>Bahia | Backcross F1 x HH<br>(hatchling) | Loggerhead<br>Rookery | 0.959 | 137.1 | 0 | 0.272 |
| Olive ridley | Hybrid Olive ridley x<br>Loggerhead | Loggerhead<br>Rookery | 0.977 | 151.8 | 0 | 0.503 |

### 3.2. Migratory patterns

We retrieved data for 15 samples with telemetry data. Our map (Figure 5) shows the post-nesting migration of loggerheads (n=7) are towards more northern sites along the Brazilian coast. Most loggerheads migrated to feeding aggregations near the coast of Ceará state (n=5), up to the Pará coast (n=2). Hawksbills migrated both north (n=3) and south (n=1) of the transmitter deployment site in Praia do Forte, with one individual migrating as far north as Ceará (Figure 5) Most hybrids exhibited a migratory pattern of loggerheads (migrating to northern areas of Pará coast) (n=3) while one hybrid migrated south of Praia do Forte.

**Figure 5.**
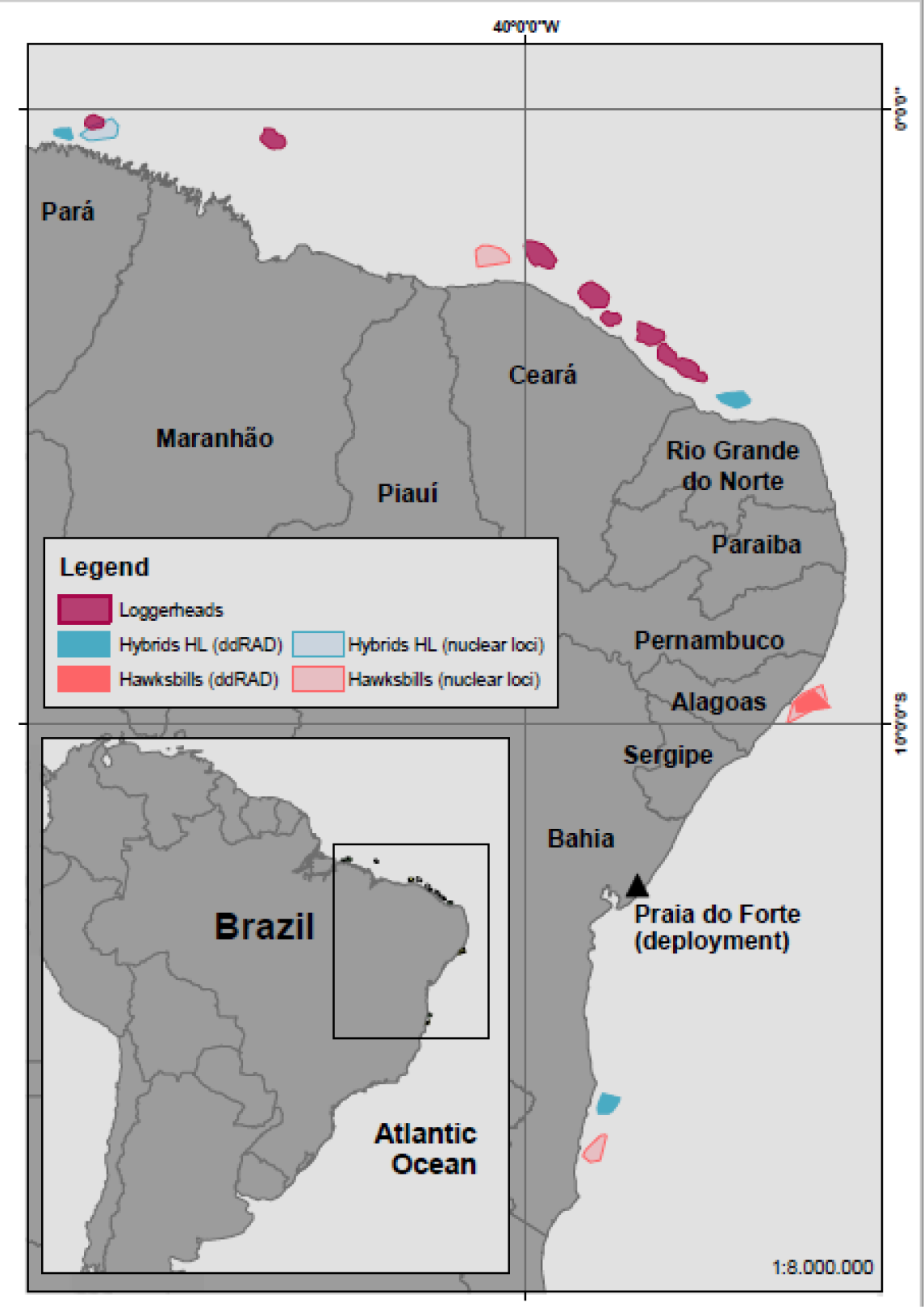
Migratory patterns for hawksbills x loggerheads (HL) hybrids (n=4), and non-hybrids hawksbill (n=4) and loggerhead (n=7) sea turtles. Specific types of genetic data (ddRAD or nuclear markers) are represented by different filling of polygons. Polygon area denotes individual interesting areas, as reported in Marcovaldi et al., (2010, 2012). Loggerheads did not have confirmed genetic data information by multiple loci, although their migration pattern was analysed and confirmed as loggerheads in Soares et al., (2021). The triangle represents Praia do Forte where samples were collected, and transmitters deployed. Modified from Marcovaldi et al., (2010, 2012).

Among the individuals with telemetry data, we obtained ddRAD data for 1 hawksbill and 3 hybrids. The remaining had only nuclear and mtDNA information. Our genomic data confirmed that three hybrids from Marcovaldi et al., (2012) were all F1 (Figure 5). The hybrid with nuclear genetic information, but no ddRAD data, exhibited a migratory pattern of loggerheads (migrating to northern areas of Pará coast). Furthermore, the individuals assigned as pure did not show indication of introgression in any molecular marker.

### 3.3. Mitogenomes

Of the 37 samples from three Brazilian rookeries and feeding sites, we obtained mitogenomes with less than 5% missing data for 15 loggerheads and 6 hawksbills. All hybrids had a loggerhead mitogenome. The remaining genomes failed to amplify at least one fragment or had low coverage regions where variants could not be called. Therefore, we considered for our analysis the 21 mitogenomes with fewer missing data and D-loop haplotypes typical of the Brazilian coast. We found a new D-loop haplotype for loggerheads (Cc-A4.4) which is related to frequent haplotypes present in the Brazilian population (Cc-A4). We also found one polymorphism (A/G) in the ND6 gene that was fixed between the two loggerhead haplotypes common in the Atlantic Ocean (Cc-A4.1 and Cc-A4.2).

### 4.4. Paternal leakage

Based on our mitogenome sequences, we successfully developed two primer pairs that specifically amplify loggerhead and hawksbill mitochondrial haplotypes typical of the Brazilian coast. The primer detection capacity was as low as a 1:10,000 dilution (Figure S3). The mixing of two loggerhead and hawksbill samples amplified both mitochondria in concentrations of 1:99 and 99:1, respectively. This means that even if the paternal mitochondria were present in small concentrations, we were able to detect it with our PCR assay. Olive ridley samples failed to amplify, but both green turtle samples were amplified by the hawksbill primer pair.

Most loggerhead x hawksbill hybrids (10 out of 12 hybrids) amplified for both fragments, indicating that they have mitochondrial DNA from both species (Table S1). Non-hybrid (“pure”) individuals (n=98, 46 loggerheads and 52 hawksbills) only amplified for their species-specific mitochondria. One F1 hybrid amplified only for the loggerhead fragment. The remaining loggerhead x hawksbill hybrid did not amplify either fragment even with multiple PCR trials. The olive ridley x loggerhead hybrid did not amplify. Amongst our samples not sequenced for ddRAD (n = 53) because of lower DNA quality/concentration, their amplification patterns agreed with morphology or previous identification using mtDNA and nuclear markers.

### 3.5. Ultra-deep sequencing of mtDNA paternal leakage

The relative abundance of the mapped reads mirrored the species assignment defined by ddRAD data and the PCR-assay amplifications. In hybrid individuals, the most abundant mitochondrial sequence was always the loggerhead’s (coverage = 390-1150X). This was an expected pattern since the female in hybrid crossings are loggerheads. In hybrids with heteroplasmy, the coverage for the hawksbill mitochondria varied from 3 to 490X and the abundance of the second mitochondria varied from 0.01 to 0.77 (Table S1). In non-hybrid samples, no sample reached the minimum average coverage threshold for the second (“leaked”) mitochondria (minimum 3X), and therefore they were all considered as homoplasmic. The backcross hatchling had no detectable second mtDNA, although its mother had the second mtDNA with 0.01 abundance. Regarding the presence of NUMTS, no regions similar to the hawksbill mtDNA were recovered in the loggerhead draft genome. All mitochondrial sequences had an identical sequence to previously reported mtDNA. We also did not detect frameshift mutations or early stop codons in protein-coding genes that could indicate possible NUMTs. All “leaked” hawksbill mitochondria had an identical sequence for the amplified fragment to other individuals found in the Bahia population. We also did not observe any polymorphisms in the sequenced region that would indicate heteroplasmy (i.e., presence of more than one type of species-specific mtDNA) besides the co-occurrence of two mtDNA types (i.e., paternal leakage) associated with two different species.

## 4. DISCUSSION

Using genomic markers, here we investigate the natural hybridization between two globally distributed sea turtle species that diverged ∼18 million years ago and are still capable of hybridizing in Brazil, and do so frequently. We showed that all adult hybrids are F1 with migration patterns more similar to loggerheads. We also showed that mitochondrial paternal leakage is present in these divergent hybrids.

### 4.1. Hybridization

Sea turtle species are separated by several million years (Vilaça et al., 2021), but have highly syntenic genomes and conserved karyotypic organization, including all species that nest in Praia do Forte (Machado et al., 2020). This may explain why hybridization could occur more frequently among sea turtles, since signatures of ancient hybridization are still found in their genomes (Vilaça et al., 2021), and may be indicate why they can still hybridize (Brito et al., 2020). Using thousands of genomic markers typed in new samples from Brazilian nesting populations and in samples collected previously (Vilaça et al., 2012), we showed that hybrid individuals from Praia do Forte (Bahia) are classified as F1s, and these were all nesting females. The only exception was a single backcross hatchling (F1 x hawksbill backcross) from a known F1 mother. Other populations along the Brazilian coast and continental shelf still did not show evidence of recent hybridization even with genome-wide markers. Furthermore, using D-statistics and *f*4-ratio we show that this is a local phenomenon in Bahia state.

Previous data from the same population showed no difference in fitness or hatchling viability between hybrids and non-hybrids of both parental species (Soares et al., 2018, 2017), but we did not find later generation hybrids among adult females. This result can be simply explained assuming that the hybridization process we observe started only few decades ago. Considering that sea turtles take 20-40 years to reach sexual maturity (Vilaça et al., 2021), >F1 adult hybrids would not be expected if hybridization started in the last 40-80 years. Conversely, if this is an older phenomenon, the lack of adults >F1 could be due to hybrid breakdown in the backcrossed hatchlings because of genomic incompatibility.

Genetic studies in the Praia do Forte population have been done since the 1980s and the presence of hybrids has been reported in all studies of this rookery (Arantes, Vilaça, et al., 2020; Bass et al., 1996; Brito et al., 2020; Conceição, Levy, Marins, & Marcovaldi, 1990; Karl et al., 1995; Lara-Ruiz et al., 2006; Proietti et al., 2014; Soares et al., 2018, 2017; Vilaça et al., 2013, 2012). However, no adult backcross has been reported yet, indicating that there is evidence of strong hybrid breakdown in the second generation. This is further supported by our RAD results, increasing marker resolution still finds all nesting females are F1s or pure species. Given the lack of >F1 adults, the time passed since the first studies detected hybrids in the 1980s, combined with similar reproductive output of F1 and backcross hybrids compared to non-hybrid individuals, we expect that F2s (F1 x F1) and backcrosses have lower or much lower fitness related to genetic incompatibilities, with no or very rare opportunities to reach sexual maturity. A similar result of lack of >F1 adult hawksbill x loggerhead hybrids was also found in a smaller nesting population in Abrolhos Archipelago (∼600 km south from Praia do Forte) where it was suggested a lower reproductive success among hatchlings of F1 hybrids when compared to loggerheads (Arantes, Ferreira, et al., 2020). In general, data from whole genomes indicated that ancient hybridization occurred in sea turtles (Vilaça et al., 2021) and left a signature in the modern genomes. Such evidence is absent in the ddRAD data, possibly as a consequence of the much lower fraction of the genome analysed by this method. Monitoring this population for the next decades, using also a reduced number of genome-wide molecular markers, is necessary to understand if the current hybridization process will produce trans-generation genomic admixture or not.

Although we did not sample male sea turtles, the use of nuclear markers can indirectly recover the paternal perspective of the hybridization. Although juvenile male loggerhead x olive ridley hybrids were found in the Brazilian coast (Medeiros et al., 2019), we have no indication that male loggerhead x hawksbill hybrids are fertile, as no F2s (F1xF1, as detected by NewHybrids) or backcrosses with male hybrids were found. Based on mtDNA and one nuclear marker sequences for >4800 hatchlings, Soares et al., (2018) observed multiple paternity patterns that could indicate male hybrids are fertile and are capable of siring offspring. Despite the large number of hatchlings, the information from one nuclear marker is not sufficient to confidently distinguish between multiple paternity involving males of multiple-species x male hybrid mating. Thousands of ddRAD loci bring important information to the hybridization dynamics by classifying the individuals as hybrids (and their hybrid generation) and also provide a higher confidence in estimating hybrid generation than few nuclear markers. Future studies using genomic markers in hatchlings or adult males could elucidate if males are indeed capable of reproduce and generate viable offspring.

### 4.2. Migratory patterns

Here we show using migratory patterns between nesting and feeding areas that 3 F1 hybrids migrated to regions known to be loggerheads’ feeding aggregations and 1 F1 hybrid migrated to a hawksbill’s aggregation. Although we hypothesized that hybrids with a migratory pattern like one species might have a larger genomic component of that species (i.e., backcrosses would display a migration pattern similar to the species’ having higher genomic proportion), here we show that such migratory pattern occurs instead in F1 hybrids. Sea turtles are dependent of migration between feeding and nesting aggregations, and it has been hypothesized that they will follow migratory trajectories that they were exposed as hatchlings (i.e., natal homing imprinting) (Brothers & Lohmann, 2015). Although no genetic components have yet been associated with migratory trajectories, migration directionality may either have a dominance type of inheritance, or it depends on oceanic currents combined with natal homing imprinting (Lohmann, Putman, & Lohmann, 2008; Mansfield et al., 2017).

Previous studies have found hybrid juveniles in hawksbills’ feeding areas in southern Brazil, and dispersal simulations showed that hybrids can potentially migrate following both species patterns (Brito et al., 2020; Proietti et al., 2014). Sea turtles are one of the classical examples of species that navigate long distances by sensing Earth’s geomagnetic field (Lohmann, Goforth, Mackiewicz, Lim, & Lohmann, 2022; Lohmann et al., 2008). Studies have shown that specific genes are involved in animal magnetoreception and might guide long-distance migration (Wan, Hayden, Iiams, & Merlin, 2021), and migration routes may also be genetically determined (Gu et al., 2021). Genomic studies of hybrids coupled with satellite tracking might help elucidate if allelic differences in migratory-related genes can explain the differential migration trajectories in hybrids.

### 4.3. Paternal leakage

Mitochondrial DNA inheritance in vertebrates is considered to be strictly maternal in most animals, although exceptions are found in various taxa (Breton & Stewart, 2015)., These unusual patterns have been associated with hybridization events as a consequence of the disruption of the cytonuclear mechanisms that ensures mtDNA’s strict maternal inheritance (as discussed below).Here we report that most fertile sea turtle hybrids have mtDNA from both parental species as a result of paternal leakage. We also show that the most abundant mtDNA is from the mother, loggerheads in all our hybrids, while the hawksbill paternal mtDNA has a lower abundance. This also confirms the directionally mating crosses in the Brazilian population.

The occurrence of mitochondrial paternal leakage happens when the sperm’s mitochondria are not eliminated during fertilization, and thus the zygote has both maternal and paternal mitochondria in its cells (Figure S4). Strict mitochondrial inheritance is thought to be controlled by ubiquitin-proteasome cellular machinery. During fertilization, the sperm’s mitochondria are tagged with ubiquitin (Rokas, Ladoukakis, & Zouros, 2003) and degraded (autophagy) in the proteasome (Song, Ballard, Yi, & Sutovsky, 2014). The recognition of the mitochondria is thought to be species-specific (Rokas et al., 2003). Therefore, when two divergent genomes are present in the same zygote as a consequence of hybridization, it is not surprising that the species-specific machinery that recognizes the mitochondria fails to degrade the paternal mitochondria. The two species that we investigated diverged ∼18 million years ago (Vilaça et al., 2021) and although they have highly syntenic chromosomes, their mitogenome is divergent (∼7% divergence) and each species has their characteristic mitochondrial lineages (Dutton et al., 2013; Jensen et al., 2019; Shamblin et al., 2014; Vargas et al., 2016; Vilaça, et al., 2022). Thus, it is expected that within-species mechanisms of paternal mitochondrial recognition and elimination are effective within sea turtle species, but not when both genomes are present (as in F1 hybrids). Hawksbill x loggerhead hybrids showed paternal leakage by both sequencing and the PCR-assay, demonstrating that paternal leakage was often associated with hybridization in sea turtles.

Although the majority of hybrids showed a pattern of paternal leakage, one F1 hybrid only showed loggerhead mtDNA. While our experiments based on newly developed species-specific PCR assay and ultra-deep sequencing can detect low abundance of paternal leakage, we cannot exclude that this hybrid has an abundance of the second mtDNA below our detection threshold. Another possibility is that F1 hybrids may still be viable with one species mtDNA (i.e., no paternal leakage) due to compensatory mechanisms, but backcrosses are not due to a higher imbalance in mitochondrial/nuclear proportions.

The backcrossed sample sequenced in our study (the offspring of a F1 female and hawksbill male) showed no detectable transmission of paternal mitochondria. The lack of leaked mtDNA is consistent with no transmission of the hawksbill mtDNA through the egg of the F1 female, and may indicate that paternal leakage does not happen in backcross mating. Another possibility is that the mother of this hatchling, sequenced in this study and which had the second mitochondria in low abundance (0.01), did not pass on the second mitochondria. Although we did not find evidence of leaked mtDNA in the backcross, we cannot discard the possibility that hawksbill mtDNA is present below our detection threshold. Thus, future studies are needed to understand the dynamics of mtDNA leakage in sea turtle hybrids. Nonetheless, the lack or extremely low abundance of the hawksbill mtDNA may be a contributing factor to why hybrid breakdown is observed in backcrosses. Since backcrossed hatchlings are viable (Soares et al., 2018), the fitness costs may become apparent in a different life stage and this might explain why no backcrosses were found in the adult population. Disruption in F2 hybrids has been associated with cytonuclear coadapted genes in other animal hybrids (Barreto et al., 2018; Han & Barreto, 2021). Higher mitonuclear fitness, as measured by oxidative phosphorylation (OXPHOS) ATP synthesis capacity, was observed in hybrid crustaceans when nuclear and mitochondrial genes were from the same parental population suggesting cytonuclear adaptions (Han & Barreto, 2021). Sea turtles’ mtDNA shows adaptive molecular evolution in genes involved in OXPHOS possibly as a response to active behavior (Ramos, Freitas, & Nery, 2020), and the higher presence of loggerhead mtDNA in backcrosses with 75% of Hawksbill genome may reduce ATP synthesis capacity and cause hybrid breakdown due to incompatible cytonuclear combinations. Studies using a larger sample set of F1 and backcrossed hatchlings might elucidate if paternal leakage plays a role in fitness and hybrid breakdown.

Our PCR assay is a possible cost-effective screening method for hybrids in the Bahia population. Our assay was designed specifically for Brazilian samples with mitochondrial haplotypes characteristic of the southwest Atlantic Ocean. Given that few mitogenomes are currently available for sea turtles, it is unclear if our assay will amplify or remain species-specific when used in individuals with haplotypes that are not present in the Brazilian loggerhead and hawksbill nesting population. An indication was the failed mitochondrial amplification in some individuals in both long-range mitogenome PCRs and our assay. The sequencing of mitogenomes from more individuals and populations outside the Atlantic Ocean will allow the design of species-specific primers capable of detecting paternal leakage in all sea turtle populations.

### 4.4. Hybrids and conservation

Sea turtles are species of conservation concern and hybridization in Brazil’s biggest nesting population needs to be better understood. Our results show that although hybrids at a first glance might represent a waste of reproductive effort, since no >F1 adult females are observed, this is a natural process and might have long-term evolutionary importance. Despite the frequent hybridization, we show evidence that there is a strong hybrid breakdown in the second generation and that the Brazilian population is not a hybrid swarm. Consequently, these two species continue to maintain distinct genetic pools and separate evolutionary trajectories. Interspecies hybridization can increase genetic diversity (Ottenburghs, 2021) and exchange adaptive alleles (Edelman & Mallet, 2021). Therefore, our recommendation is to continue the recovery efforts of the Bahia population. We expect that with the growing availability of mating partners, the presence of hybrids in Bahia population should decrease in the next decades. However, climate change is predicted to cause reproductive alterations, changes in species ranges, and skewed sex ratios (Jensen et al., 2018; Lockley & Eizaguirre, 2021), which may increase opportunities of interspecific mating in other populations beyond the Brazilian. Knowing the causes and consequences of hybridization is therefore important to understand how climate change may alter mating opportunities. Nonetheless, because of the uniqueness of this phenomenon, being one of the oldest known lineages capable of natural hybridization in the animal kingdom, they might help answer many pending questions in sea turtle biology and evolution.

## ACKNOWLEDGEMENTS

This research was supported by an Alexander von Humboldt research fellowship awarded to STV and a CNRS grant “Incubator for interdisciplinary research projects in French Guyana” awarded to DC. STV is currently supported by a European Union’s Horizon 2020 Research and Innovation Programme under the Marie Skłodowska-Curie grant agreement 844756 (TurtleHyb). We thank the BeGenDiv team for help with laboratory and bioinformatic analysis, especially Susan Mbedi and Kirsten Richter for laboratory support. Samples were collected/transported under SISBIO permit 37499-2. Tissue samples were exported from Brazil under CITES permits 13BR011173/DF and 14BR015253/DF and imported into Germany under CITES permits E-013161/13 and E-03346/14, respectively. We thank Alonzo Alfaro-Núñez for sharing primer sequences. We thank Fundação Projeto Tamar (SISBIO permit N. 42760) for logistical support in sample collection and contributions to this manuscript.

## DATA ACCESSIBILITY

All data is available on NCBI Sequence Read Archive (SRA) database under BioProject PRJNA857276. All scripts used in the ddRAD analysis are available at https://github.com/mazzoniizw/mazzoni-begendiv/. Other relevant data are available on Zenodo DOI: 10.5281/zenodo.7038996.

## AUTHOR CONTRIBUTIONS

Experimental design was performed by STV and CJM, STV performed all laboratory work, STV FM analysed the data, PL analysed migratrion data, LSA performed ddRAD data pre-processing, BdT DC FRS provided samples, GB helped interpret data, STV wrote the manuscript with input from all authors.

