## Supplementary Material for "Evidence of backcross inviability and mitochondrial DNA paternal leakage in sea turtle hybrids"

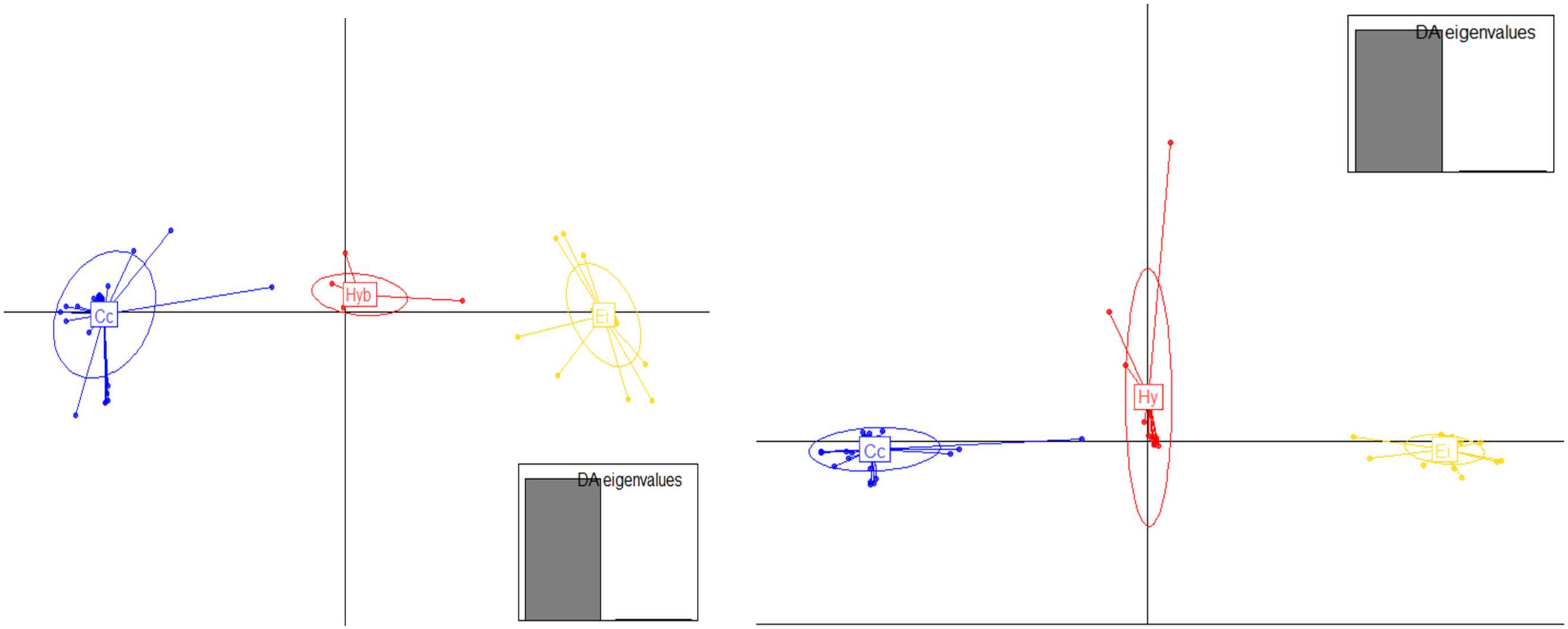

Figure S1 – DAPC plots. Left: SNPs, Right: Microhaplotypes. Cc=loggerhead, Ei=hawksbill.

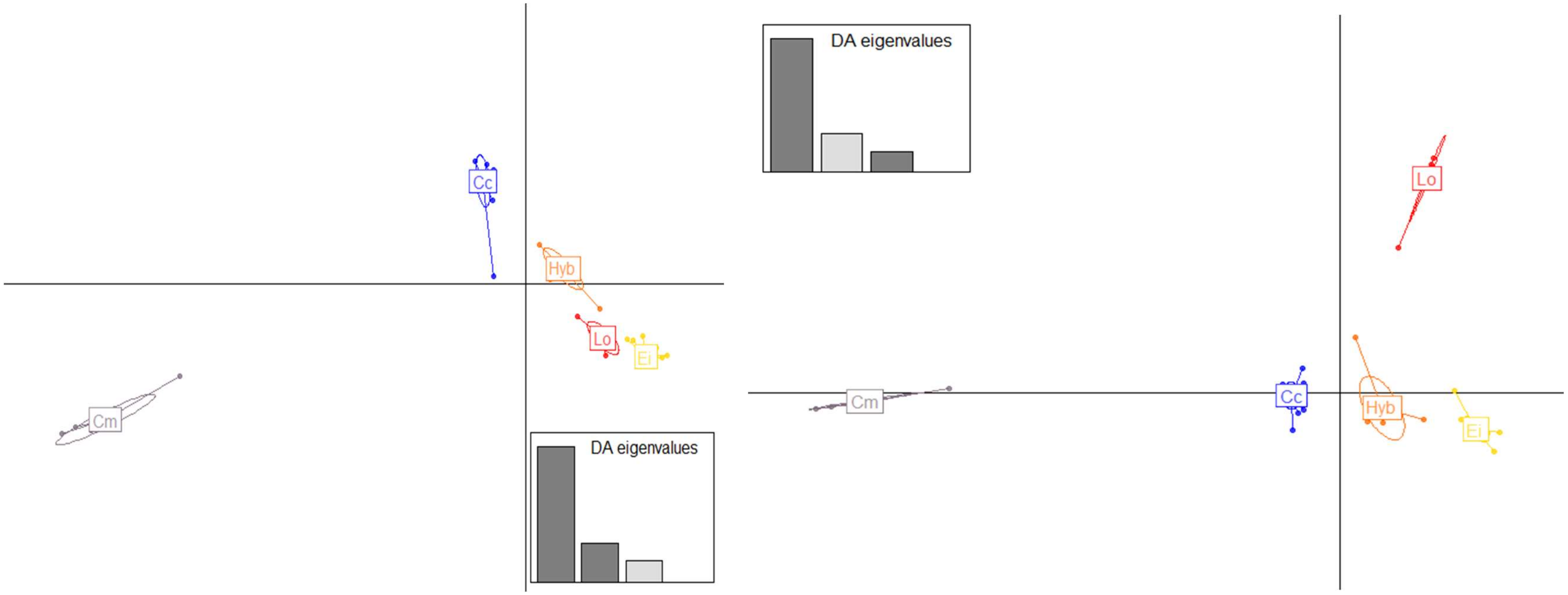

Fig S2. DAPC for the four species and their hybrids. Three Discriminant Axis were retained. First and second axis are plotted in the first graph (left); first and third axis are plotted in the second graph (right). Cc=loggerhead, Ei=hawksbill, Lo= olive ridley, Cm= green sea turtle.

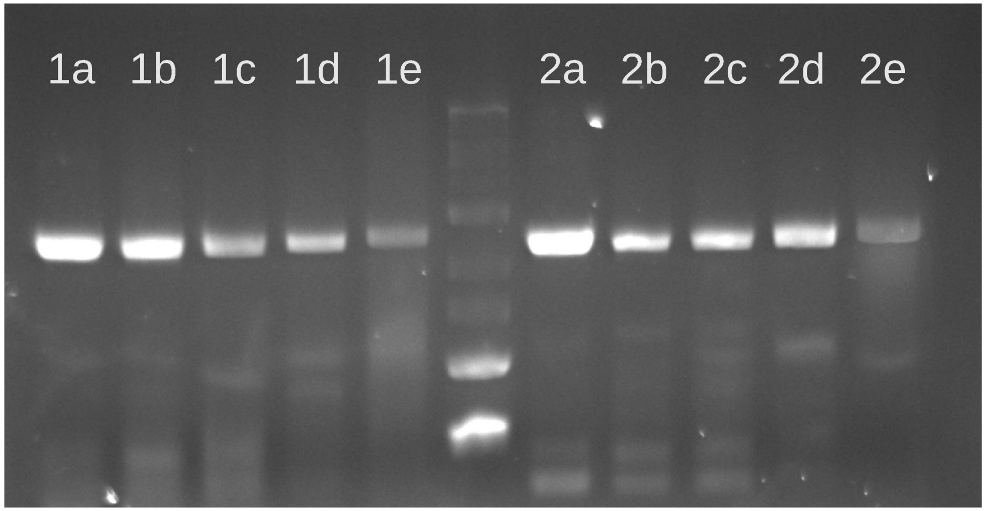

Figure S3 – Detection capacity of each primer pair by serial diluting one species DNA. Mitochondrial paternal leakage was simulated by mixing the two samples in different concentrations. Number 1 represents diluted Loggerhead samples (i.e., “leaked” loggerhead mtDNA), while 2 represents hawksbill. A) 1:1 (no dilution, 50% DNA of each species). B) 1:10. C) 1:100. D) 1:1000. E) 1:10,000.

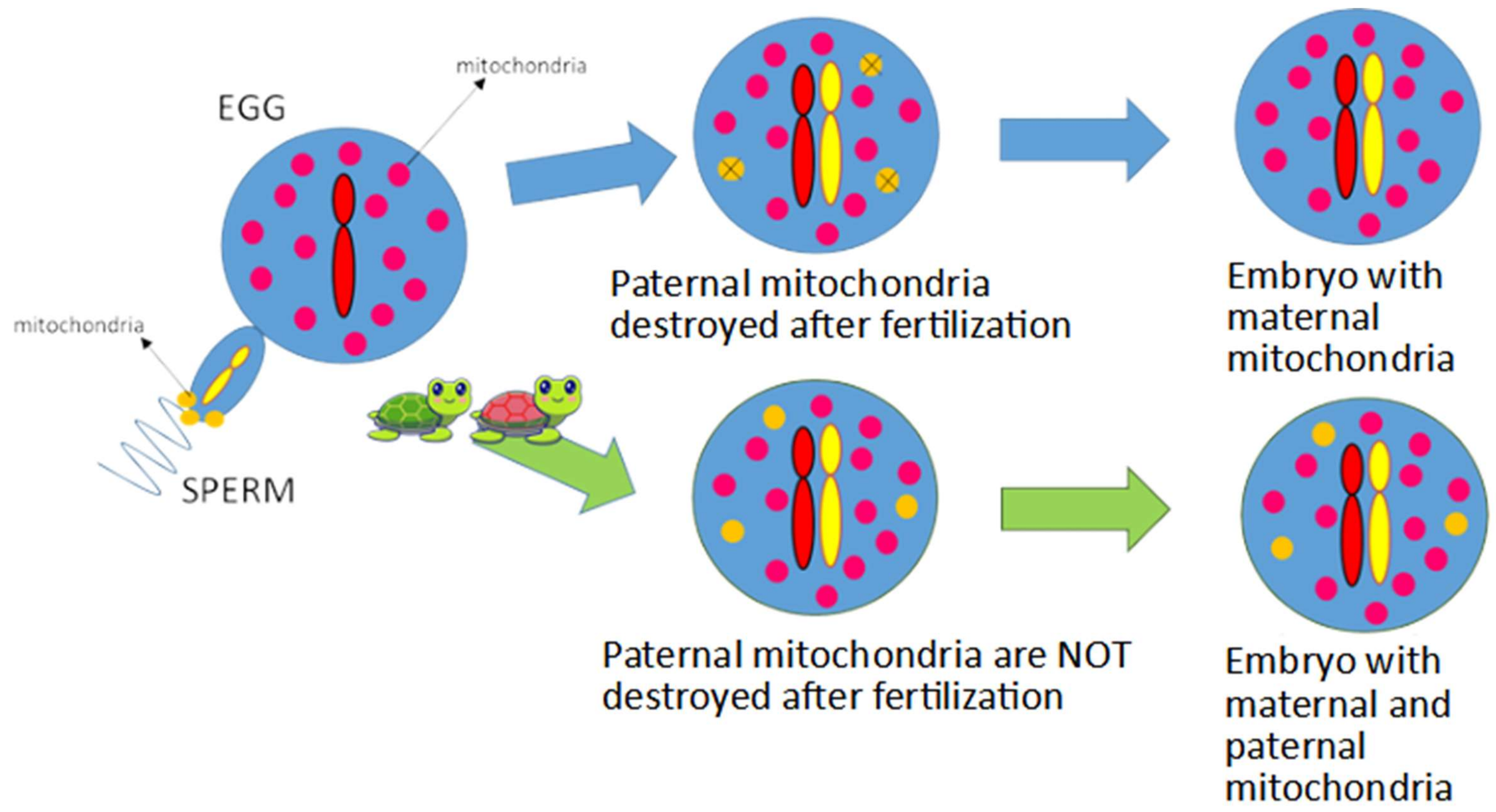

Figure S4– Scheme depicting the strict mitochondrial maternal inheritance present in most animal species (above, blue arrows), and the paternal leakage present in sea turtles (below, green arrows).

Table S1 Primer sequences and fragment sizes used for amplifications. Fragment sizes are based on the green turtle mitogenome.

| Primer F (Sequence 5′→3′) | Reference | Primer R (Sequence 5′→3′) | Reference | Fragment size (bp) | Annealing temperature (°C) |
|---|---|---|---|---|---|
| 15kbF (GGAGCCGGTATCAGGCACACC) | Duchene et al. 2012 | 6134R (GTCTCCTCCCCCTGAAGGAT) | This study | 5920 | 57 |
| 6113F (TGATCCTTCAGGGGGAGGAG) | This study | 11216R (TTTGTGTTAGTGTTGCGGCG) | This study | 5101 | 55 |
| 9518F (ATGAGGCTCATGCTCCCCTA) | This study | 12629R (GGGCCTTCTATAGCTGCTGG) | This study | 3093 | 57 |
| 12611F (CAGCAGCTATAGAAGGCCCC) | This study | H950 (GTCTCGGATTTAGGGGTTT) | Abreu-Gobrois et al. 2006 | 3801 | 55 |

Table S2 – Primer sequences used for species-specific mtDNA amplifications. Differences between the species-specific primers are underlined.

| Species | Primer F (Sequence 5′→3′) | Primer R (Sequence 5′→3′) |
|---|---|---|
| Loggerhead | TAGGC**TCC**AAAGCAGCCACC | CTAGGCC**AA**TTCATGC**C**AGG |
| Hawksbill | TAGGC**CTT**AAAGCAGCCACC | CTAGGCC**GG**TTCATGC**T**AGG |

**Table S3 - Details of results. Abbreviations: LL: loggerheads, HH: hawksbills, OL: olive ridley x loggerhead F1 hybrid, HL: loggerhead x hawksbill F1 hybrids, HHL: backcrossed hybrid, OO: olive ridleys, GG: green sea turtles.**

| **Country** | **Species** | **ID** | site | N reads ddRAD | species ID ddRAD | mtDNA D-loop haplotype | # runs deep sequencing | mean coverage loggerhead mtDNA | stsdev coverage loggerhead mtDNA | # reads loggerhead mtDNA | mean coverage hawksbill mtDNA | stdev coverage hawksbill mtDNA | # reads hawksbill mtDNA | ratio coverage | mitogenome sequences | mitogenome mean cov | mitogenome std dev | **PCR assay - mtDNA present** |
|---|---|---|---|---|---|---|---|---|---|---|---|---|---|---|---|---|---|---|
| Brazil | *C. caretta* | 221 | Praia do Forte | 1816859 | LL | | | | | | | | | | | | | Cc |
| Brazil | *E. imbricata* | 275 | Praia do Forte | 3141389 | HL | | | | | | | | | | | | | Cc+Ei |
| Brazil | *C. caretta* | 443 | Santo Antonio (SA2) | 2737721 | LL | Cc-A4.2 | 2x | 938.5 | 436 | 19331 | 0.3 | 0.5 | 7 | 0.00 | | | | Cc |
| Brazil | *C. caretta* | 445 | Santo Antonio (SA2) | 1917976 | LL | Cc-A4.1 | 2x | 914.2 | 412.1 | 19412 | 0.2 | 0.6 | 6 | 0.00 | 491343 | 4557.4 | 6703.9 | Cc |
| Brazil | *C. caretta* | 478 | G2 | | | | | | | | | | | | | | | Cc |
| Brazil | *C. caretta* | 520 | Praia do Forte | 1934395 | LL | Cc-A4.2 | 1x | | | | | | | | | | | Cc |
| Brazil | *C. caretta* | 539 | Imbassai (SA1) | 4453034 | LL | Cc-A4.1 | 2x | 1259 | 489 | 27190 | 0.4 | 0.6 | 9 | 0.00 | 448253 | 4253.9 | 8257.0 | Cc |
| Brazil | *C. caretta* | 553 | Santo Antonio (SA2) | 1447274 | LL | | | | | | | | | | | | | Cc |
| Brazil | *C. caretta* | 738 | Praia do Forte | | | Cc-A4.1 | | | | | | | | | | | | Cc |
| Brazil | *C. caretta* | 839 | Santo Antonio (SA2) | | | | | | | | | | | | | | | Cc |
| Brazil | *E. imbricata* | 840 | Imbassaí | 11403979 | HH | Ei10=Ei 11 | | | | | | | | | 347851 | 3209.5 | 4927.3 | |
| Brazil | *C. caretta* | 846 | Porto do Sauipe | 989987 | LL | | | | | | | | | | | | | Cc |
| Brazil | *C. caretta* | 851 | Subauma | | | Cc-A4.2 | | | | | | | | | | | | Cc |
| Brazil | *E. imbricata* | 881 | Imbassaí | | | | | | | | | | | | | | | Cc |
| Brazil | *C. caretta* | 985 | Imbassai | | | | | | | | | | | | | | | Cc |
| Brazil | *E. imbricata* | 1026 | Santo Antônio | 2104871 | HH | Ei8 | | | | | | | | | | | | Ei |
| Brazil | *C. caretta* | 1087 | Imbassai | | | Cc-A4.1 | | | | | | | | | | | | Cc |
| Brazil | *C. caretta* | 1141 | Praia do Forte | 3246953 | LL | Cc-A4.2 | 2x | 623 | 330.5 | 13850 | 2 | 1.8 | 41 | 0.00 | 440844 | 4101.1 | 4729.2 | Cc |
| Brazil | *C. caretta* | 1212 | Praia do Forte | | | | | | | | | | | | | | | Cc |
| Brazil | *E. imbricata* | 1451 | Santo Antônio | | | Ei10=Ei 11 | | | | | | | | | | | | Cc+Ei |
| Brazil | *C. caretta* | 1482 | Praia do Forte | 2454971 | LL | Cc-A4.2 | 2x | 1042 | 616.2 | 23301 | 0.5 | 0.6 | 10 | 0.00 | 332064 | 3098.4 | 6009.6 | Cc |
| Brazil | *C. caretta* | 1491 | Imbassai (SA1) | | | Cc-A4.2 | | | | | | | | | | | | Cc |
| Brazil | *E. imbricata* | 1494 | Praia do Forte | | | Ei8 | 2x | 2.2 | 1.8 | 54 | 1191.3 | 345.1 | 25117 | 0.00 | 548076 | 5274.5 | 9039.5 | Ei |
| Brazil | *E. imbricata* | 1562 | Praia do Forte | | | Ei10=Ei 11 | 2x | 2.3 | 2 | 48 | 991.3 | 395.3 | 21711 | 0.00 | 528922 | 4910.4 | 7914.6 | Ei |
| Brazil | *C. caretta* | 1564 | Praia do Forte | 4602691 | LL | Cc-A4.2 | 2x | 554.7 | 270.6 | 11770 | 0.9 | 1.1 | 18 | 0.00 | 461115 | 4179.8 | 5456.4 | Cc |
| Brazil | *C. caretta* | 1591 | Praia do Forte | 3527878 | LL | Cc-A4.2 | 2x | 938.5 | 380.9 | 20838 | 0.1 | 0.2 | 2 | 0.00 | 342692 | 3164.4 | 4707.0 | Cc |

| | | | | | | | | | | | | | | | | | | |
|---|---|---|---|---|---|---|---|---|---|---|---|---|---|---|---|---|---|---|
| Brazil | *C. caretta* | 1592 | Praia do Forte | 1885460 | LL | Cc-A4.2 | 2x | 1082.6 | 458.5 | 22596 | 0.5 | 0.7 | 11 | 0.00 | | | | Cc |
| Brazil | *C. caretta* | 1612 | Praia do Forte | 2663165 | LL | Cc-A4.2 | | | | | | | | | | | | Cc |
| Brazil | *E. imbricata* | 1623 | Busca Vida | | | Cc-A4.2 | 2x | 516.4 | 233.7 | 11456 | 11.7 | 7.9 | 247 | 0.02 | 87492 | 709.3 | 709.9 | |
| Brazil | *C. caretta* | 1629 | Praia do Forte | 1723014 | OL | Lo67 | | | | | | | | | | | | |
| Brazil | *C. caretta* | 1630 | Praia do Forte | 2596708 | LL | Cc-A4.2 | 2x | 1156.6 | 566.8 | 24822 | 1.5 | 1.4 | 32 | 0.00 | 391592 | 3768.0 | 7080.9 | Cc |
| Brazil | *E. imbricata* | 1637 | Praia do Forte | | | Ei8 | 2x | 4.4 | 2.6 | 93 | 1033.3 | 415 | 22418 | 0.00 | 397367 | 3747.8 | 6128.5 | Ei |
| Brazil | *E. imbricata* | 1641 | Arembepe BA | 2827680 | HL | Cc-A4.2 | 2x | 638.2 | 310.7 | 13184 | 490.4 | 226.5 | 10381 | 0.77 | 204067 | 1938.3 | 4269.9 | |
| Brazil | *E. imbricata* | 1648 | Praia do Forte | 2369833 | HL | Cc-A4.1 | 2x | 875.2 | 476.9 | 19494 | 25.5 | 12 | 536 | 0.03 | 348264 | 3172.4 | 5081.5 | Cc+Ei |
| Brazil | *C. caretta* | 1660 | Praia do Forte | | | | | | | | | | | | | | | Cc |
| Brazil | *C. caretta* | 1679 | Praia do Forte | | | | | | | | | | | | | | | Cc |
| Brazil | *C. caretta* | 1704 | Praia do Forte | 3348087 | LL | Cc-A4.1 | 2x | 973.3 | 572.8 | 21726 | 0.5 | 0.6 | 10 | 0.00 | 225765 | 2105.6 | 3769.4 | Cc |
| Brazil | *C. caretta* | 1738 | Praia do Forte | | | Cc-A4.1 | 2x | 1020.3 | 550.6 | 22479 | 1 | 10 | 22 | 0.00 | 364986 | 3473.0 | 6277.0 | Cc |
| Brazil | *E. imbricata* | 1749 | Praia do Forte | 2702059 | HH | Ei8 | 2x | 0.4 | 0.7 | 9 | 1019 | 415.5 | 22846 | 0.00 | 434927 | 3982.9 | 6848.8 | Ei |
| Brazil | *E. imbricata* | R0023 | Praia do Forte (BA) | | | CC-A4.2 | 1x | 548.7 | 254.9 | 11733 | 0.3 | 0.6 | 8 | 0.00 | 366958 | 3533.9 | 5164.7 | |
| Brazil | *E. imbricata* | R0024 | Praia do Forte (BA) | 3002068 | HL | CC-A4.1 | 1x | 548.2 | 301 | 11470 | 3.6 | 2.4 | 82 | 0.01 | | | | Cc+Ei |
| Brazil | *E. imbricata* | R0025 | Praia do Forte (BA) | 3663290 | HHL | CC-A4.1 | 1x | 432.3 | 172.5 | 9541 | 0.1 | 0.4 | 3 | 0.00 | 837701 | 7777.3 | 9580.8 | |
| Brazil | *E. imbricata* | R0040 | Praia do Forte (BA) | 3018020 | HH | EiBR8 | | | | | | | | | | | | Ei |
| Brazil | *E. imbricata* | R0053 | Praia do Forte (BA) | 2933025 | HL | CC-A4.1 | 1x | 688.5 | 333.5 | 15058 | 3 | 2.5 | 66 | 0.004 | 399496 | 3835.3 | 5481.5 | Cc+Ei |
| Brazil | *E. imbricata* | R0058 | Praia do Forte (BA) | 1987953 | HL | CC-A4.1 | | | | | | | | | | | | Cc |
| Brazil | *E. imbricata* | R0081 | Arembepe (BA) | | | EiBR8 | | | | | | | | | | | | Ei |
| Brazil | *E. imbricata* | R0089 | Arembepe (BA) | | | EiBR8 | | | | | | | | | | | | Ei |
| Brazil | *E. imbricata* | R0168 | Arembepe (BA) | | | EiBR8 | | | | | | | | | | | | Ei |
| Brazil | *E. imbricata* | R0169 | Arembepe (BA) | 2240467 | HH | EiBR8 | | | | | | | | | | | | Ei |
| Brazil | *E. imbricata* | R0186 | Arembepe (BA) | 2988207 | HL | CC-A4.2 | 1x | 676 | 302 | 14275 | 89.6 | 46.3 | 2027 | 0.13 | 296947 | 2639.5 | 3330.5 | Cc+Ei |
| Brazil | *E. imbricata* | R0187 | Arembepe (BA) | 2613289 | HL | CC-A4.2 | 1x | 544.5 | 433.6 | 12632 | 16.8 | 9.1 | 366 | 0.03 | 279314 | 2481.5 | 3581.5 | Cc+Ei |
| Brazil | *E. imbricata* | R0189 | Arembepe (BA) | 2993020 | HL | CC-A4.1 | 1x | 545.7 | 351.3 | 11778 | 8.8 | 6.4 | 186 | 0.02 | 419235 | 3832.3 | 5633.6 | Cc+Ei |
| Brazil | *E. imbricata* | R0201 | Arembepe (BA) | 2204708 | HL | CC-A4.2 | 1x | 992.7 | 535.8 | 19746 | 29.3 | 18.7 | 616 | 0.03 | 467122 | 4265.8 | 8152.0 | Cc+Ei |
| Brazil | *E. imbricata* | R0202 | Arembepe (BA) | 2938472 | HL | CC-A4.1 | 1x | 726 | 524.6 | 15716 | 20 | 18.2 | 421 | 0.03 | 442751 | 4074.2 | 6278.2 | Cc+Ei |
| Brazil | *E. imbricata* | R0218 | Praia do Forte (BA | 2360519 | HH | EiBR9 | 2x | 1031.6 | 511.4 | 22324 | 0 | 0 | 0 | 0.00 | 540226 | 5146.0 | 9106.3 | Ei |
| Brazil | *E. imbricata* | R0219 | Arembepe (BA) | 2193957 | HH | Ei10=Ei 11 | | | | | | | | | | | | |
| Rio Grande elevation | *C. caretta* | R0276 | Litoral Sul/Sudeste | 2516657 | LL | CC-A2 | 2x | 817.8 | 424.5 | 16973 | 0.1 | 0.4 | 3 | 0.00 | 190168 | 1688.3 | 2893.8 | |
| Rio Grande elevation | *C. caretta* | R0288 | Litoral Sul/Sudeste | 2193754 | LL | CC-A4 | 2x | 447.5 | 211.6 | 9826 | 0.1 | 0.4 | 4 | 0.00 | 287640 | 2570.7 | 3880.5 | Cc |
| Rio Grande elevation | *C. caretta* | R0292 | Litoral Sul/Sudeste | 2213718 | LL | CC-A2 | | | | | | | | | | | | |
| Rio Grande elevation | *C. caretta* | R0293 | Litoral Sul/Sudeste | 3721121 | LL | CC-A34 | | | | | | | | | | | | |

| | | | | | | | | | | | | | | | | | | |
|---|---|---|---|---|---|---|---|---|---|---|---|---|---|---|---|---|---|---|
| Rio Grande elevation | *C. caretta* | R0295 | Litoral Sul/Sudeste | 2168895 | LL | CC-A4 | | | | | | | | | | | | Cc |
| Rio Grande elevation | *C. caretta* | R0296 | Litoral Sul/Sudeste | | | CC-A33 | 2x | 1067.4 | 665.1 | 24311 | 1.7 | 1.4 | 32 | 0.00 | 250174 | 2276.0 | 4745.1 | |
| Rio Grande elevation | *C. caretta* | R0297 | Litoral Sul/Sudeste | 1784075 | LL | CC-A34 | | | | | | | | | | | | |
| Rio Grande elevation | *C. caretta* | R0303 | Litoral Sul/Sudeste | 2226430 | LL | CC-A4 | | | | | | | | | | | | |
| Rio Grande elevation | *C. caretta* | R0304 | Litoral Sul/Sudeste | 3165323 | LL | CC-A4 | | | | | | | | | | | | |
| Rio Grande elevation | *C. caretta* | R0306 | Litoral Sul/Sudeste | 2369414 | LL | CC-A2 | | | | | | | | | | | | |
| Rio Grande elevation | *C. caretta* | R0331 | Litoral Sul/Sudeste | 3966052 | LL | CC-A4 | | | | | | | | | | | | |
| Rio Grande elevation | *C. caretta* | 66 | | | | | | | | | | | | | | | | Cc |
| Brazil | *C. caretta* | 106 | Povoação ES | | | | | | | | | | | | | | | Cc |
| Brazil | *C. caretta* | 166 | PG | | | | | | | | | | | | | | | Cc |
| Brazil | *C. caretta* | 183 | ES | 3025645 | LL | Cc-A4.2 | | | | | | | | | | | | Cc |
| Brazil | *C. caretta* | 208 | ES | | | | | | | | | | | | | | | Cc |
| Brazil | *C. caretta* | 218 | Povoação ES | | | | | | | | | | | | | | | Cc |
| Brazil | *C. caretta* | 229 | Povoação ES | | | Cc-A4.2 | 2x | 1026.3 | 393.4 | 21350 | 0.1 | 0.2 | 1 | 0.00 | 523141 | 5037.3 | 7544.6 | |
| Brazil | *C. caretta* | 244 | ES | 3737276 | LL | | | | | | | | | | | | | Cc |
| Brazil | *C. caretta* | 278 | Povoação ES | 1810902 | LL | Cc-A4.1 | 2x | 530.4 | 437 | 10597 | 0 | 0 | 0 | 0.00 | | | | Cc |
| Brazil | *C. caretta* | 298 | Povoação ES | 2440909 | LL | | | | | | | | | | | | | Cc |
| Brazil | *C. caretta* | 323 | ES | 2331311 | LL | | | | | | | | | | | | | Cc |
| Brazil | *C. caretta* | 335 | ES | 4774186 | LL | Cc-A4.2 | | | | | | | | | | | | Cc |
| Brazil | *C. caretta* | 363 | ES | | | Cc-A4.4 | 2x | 350 | 176.3 | 7429 | 0 | 0 | 0 | 0.00 | 113101 | 915.3 | 964.9 | Cc |
| Brazil | *C. caretta* | 379 | Povoação ES | | | Cc-A4.2 | | | | | | | | | 612771 | 5795.2 | 6862.3 | Cc |
| Brazil | *C. caretta* | 417 | Povoação ES | 2355548 | LL | | | | | | | | | | | | | Cc |
| Brazil | *C. caretta* | 433 | Comboios ES | | | Cc-A4.2 | | | | | | | | | | | | Ei |
| Brazil | *C. caretta* | 561 | ES | 5544877 | LL | Cc-A4.2 | | | | | | | | | 114432 | 911.1 | 714.2 | Cc |
| Brazil | *C. caretta* | 562 | ES | 1903054 | LL | | | | | | | | | | | | | Cc |
| Brazil | *E. imbricata* | 15 | Pipa RN | 2929519 | HH | Ei8 | | | | | | | | | | | | Ei |
| Brazil | *E. imbricata* | 95 | Pipa RN | | | | | | | | | | | | | | | Ei |
| Brazil | *E. imbricata* | 135 | Pipa RN | 2718639 | HH | Ei8 | | | | | | | | | | | | Ei |
| Brazil | *E. imbricata* | 137 | Pipa RN | 2348229 | HH | Ei8 | | | | | | | | | | | | Ei |
| Brazil | *E. imbricata* | 198 | Pipa RN | | | | | | | | | | | | | | | Ei |
| Brazil | *E. imbricata* | 237 | Pipa RN | | | | | | | | | | | | | | | Ei |
| Brazil | *E. imbricata* | 239 | Pipa RN | | | | | | | | | | | | | | | Ei |
| Brazil | *E. imbricata* | 333 | Pipa RN | 2425193 | HH | Ei8 | | | | | | | | | | | | Ei |

| | | | | | | | | | | | | | | | | | | |
|---|---|---|---|---|---|---|---|---|---|---|---|---|---|---|---|---|---|---|
| Brazil | *E. imbricata* | 397 | Pipa RN | 2969470 | HH | | | | | | | | | | | | | Ei |
| Brazil | *E. imbricata* | 401 | Pipa RN | 2000703 | HH | | | | | | | | | | | | | Ei |
| Brazil | *E. imbricata* | 469 | Pipa RN | 1619619 | HH | | | | | | | | | | | | | |
| Brazil | *E. imbricata* | 472 | Pipa RN | 1746221 | HH | | | | | | | | | | | | | Ei |
| Brazil | *E. imbricata* | 473 | Pipa RN | 4333423 | HH | Ei8 | 2x | 1.6 | 1.5 | 37 | 667.4 | 230.5 | 13433 | 0.00 | 607654 | 5683.2 | 6271.5 | Ei |
| Brazil | *E. imbricata* | 475 | Pipa RN | | | | | | | | | | | | | | | Ei |
| Brazil | *E. imbricata* | 476 | Pipa RN | | | | | | | | | | | | | | | Ei |
| Brazil | *E. imbricata* | 477 | Pipa RN | | | | | | | | | | | | | | | Ei |
| Brazil | *E. imbricata* | 479 | Pipa RN | 2581770 | HH | | | | | | | | | | | | | Ei |
| Brazil | *E. imbricata* | 480 | Pipa RN | | | | | | | | | | | | | | | Ei |
| Brazil | *E. imbricata* | 482 | Pipa RN | 1172651 | HH | | | | | | | | | | | | | Ei |
| Brazil | *E. imbricata* | 488 | Pipa RN | | | | | | | | | | | | | | | Ei |
| Brazil | *E. imbricata* | 497 | Pipa RN | | | | | | | | | | | | | | | Ei |
| Brazil | *E. imbricata* | 505 | Pipa RN | | | | | | | | | | | | | | | Ei |
| Brazil | *E. imbricata* | 513 | Pipa RN | 3704874 | HH | | | | | | | | | | | | | Ei |
| Brazil | *E. imbricata* | 526 | Pipa RN | 1999420 | HH | | | | | | | | | | | | | Ei |
| Brazil | *E. imbricata* | 654 | Pipa RN | 2994328 | HH | | | | | | | | | | | | | Ei |
| Brazil | *E. imbricata* | 826 | Pipa RN | | | Ei8 | | | | | | | | | | | | Ei |
| Brazil | *E. imbricata* | 996 | Pipa RN | | | | | | | | | | | | | | | Ei |
| Brazil | *E. imbricata* | R0155 | Natal (RN) | 3659234 | HH | EiBR8 | | | | | | | | | | | | Ei |
| Brazil | *E. imbricata* | R0166 | Natal (RN) | | | EiBR8 | | | | | | | | | | | | Ei |
| Rocas Atoll | *E. imbricata* | R0027 | Atol das Rocas (RN) | | | EiBR8 | | | | | | | | | | | | Ei |
| Rocas Atoll | *E. imbricata* | R0029 | Rocas Atoll (RN) | 3608888 | HH | EiBR6 | 1x | 1.8 | 1.4 | 39 | 534.7 | 186.5 | 11539 | 0.00 | 636969 | 5650.4 | 7758.3 | Ei |
| Rocas Atoll | *E. imbricata* | R0031 | Rocas Atoll (RN) | 3751555 | HH | EiBR8 | | | | | | | | | | | | Ei |
| Rocas Atoll | *E. imbricata* | R0032 | Rocas Atoll (RN) | 1943196 | HH | EiBR8 | 1x | 0.4 | 0.7 | 10 | 589.5 | 224.4 | 12389 | 0.00 | 524365 | 4841.0 | 5882.9 | Ei |
| Rocas Atoll | *E. imbricata* | R0033 | Rocas Atoll (RN) | 1311475 | HH | EiBR8 | | | | | | | | | | | | Ei |
| Rocas Atoll | *E. imbricata* | R0034 | Rocas Atoll (RN) | 1293576 | HH | EiBR8 | | | | | | | | | | | | Ei |
| Rocas Atoll | *E. imbricata* | R0035 | Rocas Atoll (RN) | 3726229 | HH | EiBR12 | 1x | 670.3 | 423.6 | 14667 | 0.3 | 0.6 | 8 | 0.00 | 353594 | 3233.6 | 4511.6 | Ei |
| Rocas Atoll | *E. imbricata* | R0038 | Rocas Atoll (RN) | 3581309 | HH | EiBR8 | | | | | | | | | | | | Ei |
| Rocas Atoll | *E. imbricata* | R0039 | Rocas Atoll (RN) | 1152296 | HH | EiBR8 | | | | | | | | | | | | Ei |
| Rocas Atoll | *E. imbricata* | R0068 | Rocas Atoll (RN) | 1634466 | HH | EiBR5 | | | | | | | | | | | | Ei |
| Rocas Atoll | *E. imbricata* | R0071 | Rocas Atoll (RN) | 1349393 | HH | EiBR7 | | | | | | | | | | | | Ei |
| Rocas Atoll | *E. imbricata* | R0241 | Atol das Rocas (RN) | | | EiBR8 | | | | | | | | | | | | Ei |
| Brazil | *E. imbricata* | R0141 | Pirambu (SE) | | | Cc-A4.2 | | | | | | | | | | | | Cc+Ei |
| Brazil | *E. imbricata* | R0154 | Pirambu (SE) | | | Cc-A4.1 | | | | | | | | | | | | Cc+Ei |

| | | | | | |
|---|---|---|---|---|---|
| Brazil | *C. mydas* | 8 | | 1141102 | *GG* |
| Brazil | *C. mydas* | 49 | | 1030021 | *GG* |
| Brazil | *C. mydas* | 66 | | 882979 | *GG* |
| Brazil | *C. mydas* | 96 | | 2870709 | *GG* |
| Guadaloupe | *C. mydas* | R340 | | 5279324 | *GG* |
| Guyane | *C. mydas* | CM2 | | 4652309 | *GG* |
| French Guyane | *C. mydas* | Ei01 | | 2106684 | *GG* |
| Martinique | *C. mydas* | R375 | | 5955444 | *GG* |
| Brazil | *L. olivacea* | 474 | | 1376289 | *OO* |
| Brazil | *L. olivacea* | 1524 | | 1818400 | *OO* |
| Guyane | *L. olivacea* | LO2 | | 1525523 | *OO* |
| Guyane | *L. olivacea* | LO260 | | 1170611 | *OO* |
| Guyane | *L. olivacea* | LO3 | | 1480923 | *OO* |
| Guyane | *L. olivacea* | LO39 | | 2526487 | *OO* |
| Guyane | *L. olivacea* | LO65 | | 1207309 | *OO* |

**Table S4. Number of reads for ddRAD data after each preprocessing step. Percentages (%) refer to the previous step.**

| Sample | Species | Location | Demultiplexed reads | After Short Reads Filter | % | After Quality (Q>25) and Size Filter | % | After Check MseI | % | After Restriction Sites Filter | % |
|---|---|---|---|---|---|---|---|---|---|---|---|
| 221 | *C. caretta* | Bahia (BR) | 1816859 | 1223349 | 67.3 | 1173174 | 95.9 | 1168167 | 99.6 | 1108266 | 94.9 |
| 443 | *C. caretta* | Bahia (Brazil) | 2737721 | 2298183 | 83.9 | 2156977 | 93.9 | 2139874 | 99.2 | 1896985 | 88.6 |
| 445 | *C. caretta* | Bahia (Brazil) | 1917976 | 504218 | 26.3 | 446262 | 88.5 | 443421 | 99.4 | 426586 | 96.2 |
| 520 | *C. caretta* | Bahia (Brazil) | 1934395 | 1056347 | 54.6 | 988105 | 93.5 | 984847 | 99.7 | 882586 | 89.6 |
| 539 | *C. caretta* | Bahia (Brazil) | 4453034 | 3989940 | 89.6 | 3796936 | 95.2 | 3758063 | 99.0 | 3588639 | 95.5 |
| 553 | *C. caretta* | Bahia (Brazil) | 1447274 | 801344 | 55.4 | 756997 | 94.5 | 749418 | 99.0 | 715142 | 95.4 |
| 846 | *C. caretta* | Bahia (Brazil) | 989987 | 565056 | 57.1 | 531169 | 94.0 | 528353 | 99.5 | 440672 | 83.4 |
| 1141 | *C. caretta* | Bahia (Brazil) | 3246953 | 2835975 | 87.3 | 2667969 | 94.1 | 2654107 | 99.5 | 2538039 | 95.6 |
| 1482 | *C. caretta* | Bahia (Brazil) | 2454971 | 1411748 | 57.5 | 1343517 | 95.2 | 1334309 | 99.3 | 1287059 | 96.5 |
| 1564 | *C. caretta* | Bahia (Brazil) | 4602691 | 4281288 | 93.0 | 4142642 | 96.8 | 4113583 | 99.3 | 3999772 | 97.2 |

| | | | | | | | | | | |
|---|---|---|---|---|---|---|---|---|---|---|
| 1591 | *C. caretta* | Bahia (Brazil) | 3527878 | 2943283 | 83.4 | 2774492 | 94.3 | 2756873 | 99.4 | 2489221 | 90.3 |
| 1592 | *C. caretta* | Bahia (Brazil) | 1885460 | 525966 | 27.9 | 462388 | 87.9 | 459487 | 99.4 | 432188 | 94.1 |
| 1612 | *C. caretta* | Bahia (Brazil) | 2663165 | 1277077 | 48.0 | 1204020 | 94.3 | 1196689 | 99.4 | 1135998 | 94.9 |
| 1630 | *C. caretta* | Bahia (Brazil) | 2596708 | 2151817 | 82.9 | 2048198 | 95.2 | 2024794 | 98.9 | 1900677 | 93.9 |
| 1704 | *C. caretta* | Bahia (Brazil) | 3348087 | 1647431 | 49.2 | 1510314 | 91.7 | 1505367 | 99.7 | 1243436 | 82.6 |
| 183 | *C. caretta* | Espírito Santo (Brazil) | 3025645 | 2356758 | 77.9 | 2202112 | 93.4 | 2188071 | 99.4 | 1823749 | 83.3 |
| 244 | *C. caretta* | Espírito Santo (Brazil) | 3737276 | 3448852 | 92.3 | 3274166 | 94.9 | 3239954 | 99.0 | 3140049 | 96.9 |
| 298 | *C. caretta* | Espírito Santo (Brazil) | 2440909 | 558797 | 22.9 | 510316 | 91.3 | 506303 | 99.2 | 493708 | 97.5 |
| 323 | *C. caretta* | Espírito Santo (Brazil) | 2331311 | 906545 | 38.9 | 851210 | 93.9 | 844460 | 99.2 | 821243 | 97.3 |
| 335 | *C. caretta* | Espírito Santo (Brazil) | 4774186 | 4438663 | 93.0 | 4229347 | 95.3 | 4177958 | 98.8 | 3894229 | 93.2 |
| 417 | *C. caretta* | Espírito Santo (Brazil) | 2355548 | 1988033 | 84.4 | 1891493 | 95.1 | 1873304 | 99.0 | 1796276 | 95.9 |
| 561 | *C. caretta* | Espírito Santo (Brazil) | 5544877 | 534110 | 9.6 | 343961 | 64.4 | 341159 | 99.2 | 332991 | 97.6 |
| 562 | *C. caretta* | Espírito Santo (Brazil) | 1903054 | 1416126 | 74.4 | 1357418 | 95.9 | 1350386 | 99.5 | 1312518 | 97.2 |
| 278 | *C. caretta* | Espírito Santo (Brazil) | 1810902 | 716106 | 39.5 | 659895 | 92.2 | 656063 | 99.4 | 579443 | 88.3 |
| R276 | *C. caretta* | Rio Grande Elevation (Brazil) | 2516657 | 1764341 | 70.1 | 1693236 | 96.0 | 1686087 | 99.6 | 1634214 | 96.9 |
| R288 | *C. caretta* | Rio Grande Elevation (Brazil) | 2193754 | 800657 | 36.5 | 699821 | 87.4 | 697037 | 99.6 | 581408 | 83.4 |
| R292 | *C. caretta* | Rio Grande Elevation (Brazil) | 2213718 | 621172 | 28.1 | 578017 | 93.1 | 574128 | 99.3 | 553846 | 96.5 |
| R293 | *C. caretta* | Rio Grande Elevation (Brazil) | 3721121 | 3264441 | 87.7 | 3101433 | 95.0 | 3068002 | 98.9 | 2953998 | 96.3 |
| R295 | *C. caretta* | Rio Grande Elevation (Brazil) | 2168895 | 925100 | 42.7 | 877280 | 94.8 | 870649 | 99.2 | 846855 | 97.3 |
| R297 | *C. caretta* | Rio Grande Elevation (Brazil) | 1784075 | 903855 | 50.7 | 830541 | 91.9 | 823170 | 99.1 | 711733 | 86.5 |
| R303 | *C. caretta* | Rio Grande Elevation (Brazil) | 2226430 | 885715 | 39.8 | 828473 | 93.5 | 823920 | 99.5 | 782562 | 95.0 |
| R304 | *C. caretta* | Rio Grande Elevation (Brazil) | 3165323 | 2454168 | 77.5 | 2291262 | 93.4 | 2274561 | 99.3 | 2113573 | 92.9 |
| R306 | *C. caretta* | Rio Grande Elevation (Brazil) | 2369414 | 2134316 | 90.1 | 2069213 | 96.9 | 2041650 | 98.7 | 1936260 | 94.8 |
| R331 | *C. caretta* | Rio Grande Elevation (Brazil) | 3966052 | 3084857 | 77.8 | 2991637 | 97.0 | 2970202 | 99.3 | 2857163 | 96.2 |
| R370 | *C. caretta* | Rio Grande Elevation (Brazil) | 2876336 | 2648288 | 92.1 | 2522236 | 95.2 | 2476006 | 98.2 | 1813718 | 73.3 |
| 840 | *E. imbricata* | Bahia (Brazil) | 11403979 | 4195276 | 36.8 | 3873881 | 92.3 | 3860718 | 99.7 | 3742557 | 96.9 |
| 1026 | *E. imbricata* | Bahia (Brazil) | 2104871 | 1582732 | 75.2 | 1529367 | 96.6 | 1523572 | 99.6 | 1447081 | 95.0 |
| 1749 | *E. imbricata* | Bahia (Brazil) | 2702059 | 1895658 | 70.2 | 1779381 | 93.9 | 1772339 | 99.6 | 1466804 | 82.8 |
| R169 | *E. imbricata* | Bahia (Brazil) | 2240467 | 1773804 | 79.2 | 1711184 | 96.5 | 1703100 | 99.5 | 1641374 | 96.4 |
| R218 | *E. imbricata* | Bahia (Brazil) | 2360519 | 607284 | 25.7 | 528254 | 87.0 | 525443 | 99.5 | 484679 | 92.2 |
| R219 | *E. imbricata* | Bahia (Brazil) | 2193957 | 1927748 | 87.9 | 1856035 | 96.3 | 1850202 | 99.7 | 1793994 | 97.0 |
| R40 | *E. imbricata* | Bahia (Brazil) | 3018020 | 2372771 | 78.6 | 2278311 | 96.0 | 2268167 | 99.6 | 2189967 | 96.6 |
| 15 | *E. imbricata* | Rio Grande do Norte (Brazil) | 2929519 | 2381257 | 81.3 | 2290830 | 96.2 | 2276144 | 99.4 | 2200974 | 96.7 |
| 135 | *E. imbricata* | Rio Grande do Norte (Brazil) | 2718639 | 1905407 | 70.1 | 1823132 | 95.7 | 1812558 | 99.4 | 1716386 | 94.7 |
| 137 | *E. imbricata* | Rio Grande do Norte (Brazil) | 2348229 | 1778821 | 75.8 | 1703699 | 95.8 | 1690671 | 99.2 | 1583157 | 93.6 |
| 333 | *E. imbricata* | Rio Grande do Norte (Brazil) | 2425193 | 603408 | 24.9 | 536501 | 88.9 | 533146 | 99.4 | 492880 | 92.4 |
| 397 | *E. imbricata* | Rio Grande do Norte (Brazil) | 2969470 | 1494711 | 50.3 | 1426434 | 95.4 | 1419007 | 99.5 | 1365118 | 96.2 |
| 401 | *E. imbricata* | Rio Grande do Norte (Brazil) | 2000703 | 517556 | 25.9 | 469806 | 90.8 | 466618 | 99.3 | 447749 | 96.0 |
| 469 | *E. imbricata* | Rio Grande do Norte (Brazil) | 1619619 | 476744 | 29.4 | 431125 | 90.4 | 428902 | 99.5 | 409638 | 95.5 |

| | | | | | | | | | | |
|---|---|---|---|---|---|---|---|---|---|---|
| 472 | *E. imbricata* | Rio Grande do Norte (Brazil) | 1746221 | 375573 | 21.5 | 341434 | 90.9 | 340009 | 99.6 | 328606 | 96.6 |
| 473 | *E. imbricata* | Rio Grande do Norte (Brazil) | 4333423 | 3808310 | 87.9 | 3630325 | 95.3 | 3614645 | 99.6 | 3396754 | 94.0 |
| 479 | *E. imbricata* | Rio Grande do Norte (Brazil) | 2581770 | 1716886 | 66.5 | 1649605 | 96.1 | 1640996 | 99.5 | 1598846 | 97.4 |
| 482 | *E. imbricata* | Rio Grande do Norte (Brazil) | 1172651 | 694432 | 59.2 | 667759 | 96.2 | 663809 | 99.4 | 642640 | 96.8 |
| 513 | *E. imbricata* | Rio Grande do Norte (Brazil) | 3704874 | 2879774 | 77.7 | 2767308 | 96.1 | 2753914 | 99.5 | 2637593 | 95.8 |
| 526 | *E. imbricata* | Rio Grande do Norte (Brazil) | 1999420 | 1503544 | 75.2 | 1457132 | 96.9 | 1448089 | 99.4 | 1391778 | 96.1 |
| 654 | *E. imbricata* | Rio Grande do Norte (Brazil) | 2994328 | 2783533 | 93.0 | 2671493 | 96.0 | 2651629 | 99.3 | 2536136 | 95.6 |
| R155 | *E. imbricata* | Rio Grande do Norte (Brazil) | 3659234 | 986917 | 27.0 | 878274 | 89.0 | 874295 | 99.5 | 659492 | 75.4 |
| R29 | *E. imbricata* | Rocas Atoll (Brazil) | 3608888 | 2917079 | 80.8 | 2806474 | 96.2 | 2796949 | 99.7 | 1760822 | 63.0 |
| R31 | *E. imbricata* | Rocas Atoll (Brazil) | 3751555 | 2977526 | 79.4 | 2853199 | 95.8 | 2840072 | 99.5 | 2681784 | 94.4 |
| R32 | *E. imbricata* | Rocas Atoll (Brazil) | 1943196 | 1201974 | 61.9 | 1153306 | 96.0 | 1148851 | 99.6 | 943503 | 82.1 |
| R33 | *E. imbricata* | Rocas Atoll (Brazil) | 1311475 | 779560 | 59.4 | 746631 | 95.8 | 743968 | 99.6 | 722631 | 97.1 |
| R34 | *E. imbricata* | Rocas Atoll (Brazil) | 1293576 | 766334 | 59.2 | 737418 | 96.2 | 734382 | 99.6 | 716273 | 97.5 |
| R35 | *E. imbricata* | Rocas Atoll (Brazil) | 3726229 | 2850496 | 44.2 | 2705260 | 94.9 | 2697480 | 99.7 | 2606710 | 96.6 |
| R38 | *E. imbricata* | Rocas Atoll (Brazil) | 3581309 | 3196959 | 89.3 | 3070724 | 96.1 | 3056823 | 99.5 | 2941202 | 96.2 |
| R39 | *E. imbricata* | Rocas Atoll (Brazil) | 1152296 | 524679 | 45.5 | 494610 | 94.3 | 491496 | 99.4 | 478454 | 97.3 |
| R68 | *E. imbricata* | Rocas Atoll (Brazil) | 1634466 | 1105535 | 67.6 | 1060833 | 96.0 | 1057312 | 99.7 | 1011874 | 95.7 |
| R71 | *E. imbricata* | Rocas Atoll (Brazil) | 1349393 | 757468 | 56.1 | 722829 | 95.4 | 720610 | 99.7 | 683110 | 94.8 |
| 275 | Hybrid (HL) | Bahia (Brazil) | 3141389 | 478277 | 15.2 | 411414 | 86.0 | 406527 | 98.8 | 389542 | 95.8 |
| 1629 | Hybrid (OL) | Bahia (Brazil) | 1723014 | 521222 | 30.3 | 461545 | 88.6 | 458493 | 99.3 | 416531 | 90.8 |
| 1641 | Hybrid (HL) | Bahia (Brazil) | 2827680 | 971702 | 34.4 | 895951 | 92.2 | 892834 | 99.7 | 835006 | 93.5 |
| 1648 | Hybrid (HL) | Bahia (Brazil) | 2369833 | 739863 | 31.2 | 653584 | 88.3 | 651456 | 99.7 | 604338 | 92.8 |
| R186 | Hybrid (HL) | Bahia (Brazil) | 2988207 | 1930653 | 64.6 | 1784804 | 92.4 | 1768882 | 99.1 | 1634052 | 92.4 |
| R187 | Hybrid (HL) | Bahia (Brazil) | 2613289 | 733727 | 28.1 | 623501 | 85.0 | 621262 | 99.6 | 528422 | 85.1 |
| R189 | Hybrid (HL) | Bahia (Brazil) | 2993020 | 1824768 | 61.0 | 1658700 | 90.9 | 1640016 | 98.9 | 1480619 | 90.3 |
| R201 | Hybrid (HL) | Bahia (Brazil) | 2204708 | 1999013 | 90.7 | 1916220 | 95.9 | 1909890 | 99.7 | 1527702 | 80.0 |
| R202 | Hybrid (HL) | Bahia (Brazil) | 2938472 | 2791804 | 95.0 | 2664183 | 95.4 | 2635025 | 98.9 | 1973904 | 74.9 |
| R24 | Hybrid (HL) | Bahia (Brazil) | 3002068 | 1334932 | 44.5 | 1223847 | 91.7 | 1219699 | 99.7 | 1054700 | 86.5 |
| R25 | Hybrid (HHL | Bahia (Brazil) | 3663290 | 2285786 | 62.4 | 2121913 | 92.8 | 2104084 | 99.2 | 1650978 | 78.5 |
| R53 | Hybrid (HL) | Bahia (Brazil) | 2933025 | 1015459 | 34.6 | 911338 | 89.7 | 908313 | 99.7 | 779989 | 85.9 |
| R58 | Hybrid (HL) | Bahia (Brazil) | 1987953 | 704301 | 35.4 | 654737 | 93.0 | 651880 | 99.6 | 616128 | 94.5 |
| 8 | *C. mydas* | Brazil | 1141102 | 518202 | 45.4 | 499811 | 96.5 | 496152 | 99.3 | 479777 | 96.7 |
| 49 | *C. mydas* | Brazil | 1030021 | 394467 | 38.3 | 379427 | 96.2 | 377621 | 99.5 | 354345 | 93.8 |
| 66 | *C. mydas* | Brazil | 882979 | 490055 | 55.5 | 473661 | 96.7 | 471019 | 99.4 | 450808 | 95.7 |
| 96 | *C. mydas* | Brazil | 2870709 | 2800507 | 97.6 | 2747452 | 98.1 | 2682860 | 97.6 | 1878365 | 70.0 |
| R340 | *C. mydas* | Guadeloupe | 5279324 | 5029541 | 95.3 | 4897169 | 97.4 | 4865880 | 99.4 | 4609119 | 94.7 |
| CM2 | *C. mydas* | French Guiana | 4652309 | 4335090 | 93.2 | 4226509 | 97.5 | 4203144 | 99.4 | 3272599 | 77.9 |
| Ei01 | *C. mydas* | French Guiana | 2106684 | 994856 | 47.2 | 994856 | 100.0 | 952703 | 95.8 | 927600 | 97.4 |
| R375 | *C. mydas* | Martinique | 5955444 | 5639566 | 94.7 | 5477894 | 97.1 | 5458243 | 99.6 | 4978625 | 91.2 |

| | | | | | | | | | | | |
|---|---|---|---|---|---|---|---|---|---|---|---|
| 474 | *L.olivacea* | Brazil | 1376289 | 992666 | 72.1 | 962714 | 97.0 | 951664 | 98.9 | 859962 | 90.4 |
| 1524 | *L.olivacea* | Brazil | 1818400 | 1551114 | 85.3 | 1515860 | 97.7 | 1506452 | 99.4 | 1407285 | 93.4 |
| LO2 | *L.olivacea* | French Guiana | 1525523 | 345647 | 22.7 | 316625 | 91.6 | 315090 | 99.5 | 304443 | 96.6 |
| LO260 | *L.olivacea* | French Guiana | 1170611 | 672857 | 57.5 | 648813 | 96.4 | 641090 | 98.8 | 611740 | 95.4 |
| LO3 | *L.olivacea* | French Guiana | 1480923 | 626979 | 42.3 | 597839 | 95.4 | 589272 | 98.6 | 536743 | 91.1 |
| LO39 | *L.olivacea* | French Guiana | 2526487 | 2196771 | 86.9 | 2123461 | 96.7 | 2101272 | 99.0 | 1922652 | 91.5 |
| LO65 | *L.olivacea* | French Guiana | 1207309 | 823681 | 68.2 | 799563 | 97.1 | 792796 | 99.2 | 743986 | 93.8 |
